# Too little and too much: medial prefrontal functional inhibition impairs early, whereas neural disinhibition impairs serial reversal performance in rats

**DOI:** 10.1101/2025.02.10.637529

**Authors:** Jacco G. Renström, Charlotte J.L. Taylor, Rachel Grasmeder Allen, Luke O’Hara, Joanna Loayza, Jacob Juty, Paula M. Moran, Moritz von Heimendahl, Serena Deiana, Johann Du Hoffman, Carl W. Stevenson, Silvia Maggi, Tobias Bast

**Affiliations:** School of Psychology, University of Nottingham, NG7 2RD, United Kingdom; Boehringer Ingelheim Pharma GmbH & Co. KG, 88400 Biberach, Germany; School of Biosciences, University of Nottingham, LE12 5RD, United Kingdom

**Author notes:** Corresponding author email Jacco Renström –, Tobias Bast –.

## Abstract

Schizophrenia is associated with reduced activation (‘hypofrontality’) and neural disinhibition (reduced GABAergic inhibition) in the dorsolateral prefrontal cortex (dlPFC), as well as reversal learning deficits. Whilst reversal learning has been strongly linked to the orbitofrontal cortex, its dependence on the primate dlPFC – and its rodent analogue, the medial PFC (mPFC) – is less clear. Nevertheless, we hypothesized that the mPFC may be required for reversal learning if the reversal is demanding. Furthermore, even if the mPFC is not required, mPFC disinhibition may impair reversals, because it may disrupt processing in mPFC projection sites. To test these hypotheses, we combined bi-directional manipulations of mPFC GABAergic inhibition, using intracerebral drug microinfusion and chemogenetic/DREADD methods, with reversal testing on a food-reinforced two-lever discrimination task in rats. First, we induced mPFC functional inhibition and disinhibition, by microinfusion of the GABA-A receptor agonist muscimol or antagonist picrotoxin, respectively, and examined the impact on early reversals (reversals 1-3) and well-established serial reversals (reversal 5 onwards). Using classical performance measures and Bayesian trial-by-trial strategy analysis, we found that mPFC muscimol impaired early, but not serial, reversals, increasing perseveration and impairing exploratory (lose-shift) behavior at reversal 2. In contrast, mPFC picrotoxin impaired serial reversals, reducing exploratory (lose-shift) and exploitative (win-stay) behavior. Second, to inhibit mPFC GABAergic neurons, we expressed the inhibitory DREADD hM4Di in these neurons; chemogenetic mPFC disinhibition by activation of hM4Di also impaired serial reversal learning, primarily disrupting exploitation. Our findings suggest that mPFC hypoactivation and disinhibition disrupt distinct aspects of reversal learning by different mechanisms.

**Significance statement:** Schizophrenia is associated with reduced activation (“hypofrontality”) and neural disinhibition (reduced GABAergic inhibition) within the prefrontal cortex (PFC). Yet, it is not clear if and how these distinct aspects of prefrontal dysfunction contribute to impaired reversal learning, a key feature of the cognitive inflexibility characterizing schizophrenia. Here, we combined bi-directional manipulations of prefrontal GABAergic inhibition with testing of reversal learning in rats. Increasing prefrontal functional inhibition (i.e., reducing prefrontal activation) selectively impaired early reversals, enhancing perseveration and reducing exploratory (lose-shift) behavior, whereas prefrontal disinhibition disrupted serial reversals, impairing both exploration and exploitation. Our findings suggest that reduced activation and disinhibition of PFC disrupt distinct aspects of reversal learning, by distinct mechanisms.

## INTRODUCTION

Schizophrenia has been associated with two key neural abnormalities in the dorsolateral prefrontal cortex (dlPFC): reduced activation (hypofrontality) (Carter et al., 1998; Minzenberg et al., 2009) and reduced GABAergic inhibition (neural disinhibition)(Schoonover et al., 2020). These two aspects of prefrontal dysfunction were suggested to be causally linked, whereby hyperexcitability due to neural disinhibition at early stages of schizophrenia contributes to hypofrontality at later stages (Krystal & Anticevic, 2015). Moreover, research in animal models suggests that such imbalanced prefrontal activity, both too little and too much, can cause cognitive impairments relevant to schizophrenia and other neuropsychiatric disorders (Bast et al., 2017; Enomoto et al., 2011; Pezze et al., 2014; Rao et al., 2000; Tse et al., 2015).

Schizophrenia has also been associated with reversal learning deficits (Izquierdo et al., 2017; Leeson et al., 2009; Murray et al., 2008). Reversal learning involves the reversal of response-reward associations, whereby a previously rewarded response is no longer rewarded, whereas a previously unrewarded response becomes rewarded. Reversal learning has mainly been associated with the orbitofrontal cortex (OFC), with less importance ascribed to the dlPFC in primates and the medial PFC (mPFC) in rodents (Leeson et al., 2009; Izquierdo et al., 2017), which shares functional-anatomical properties with the human dlPFC (Brown & Bowman, 2002; Dalley et al., 2004; Izquierdo, 2024; Laubach et al., 2018; Seamans et al., 2008; Uylings et al., 2003). Key evidence for this comes from lesion and pharmacological inactivation studies showing that OFC, but not dlPFC or mPFC, was required for reversal learning (Boulougouris et al., 2007; Chudasama & Robbins, 2003; Floresco et al., 2008; Hervig et al., 2020; Izquierdo et al., 2004; McAlonan & Brown, 2003; Murray et al., 2007).

Nevertheless, imbalanced mPFC activity may impair reversal learning. First, mPFC lesions impaired reversal learning on attentionally demanding reversal paradigms, requiring difficult discriminations or simultaneous monitoring of multiple stimuli (Brigman & Rothblat, 2008; Bussey et al., 1997; Chudasama & Robbins, 2003; Kosaki & Watanabe, 2012). Second, the mPFC has been implicated in overcoming prepotent behavioral responses and in attentional set formation (Haddon & Killcross, 2007; Knott et al., 2024; Marquis et al., 2007; Miller & Cohen, 2001), which may be particularly relevant at early reversal-learning stages (Guise & Shapiro, 2017; Peters et al., 2013). Third, even under conditions when the mPFC is not required, mPFC disinhibition may disrupt reversal learning by causing aberrant drive of mPFC projections and, thereby, disrupting processing in mPFC projections sites, which include the OFC (Sesack et al., 1989).

Here, we investigated how mPFC functional inhibition and disinhibition affect early and serial reversal learning in rats, using a food-reinforced two-lever reversal task (Brady & Floresco, 2015; Gonçalves et al., 2023). To manipulate mPFC GABAergic inhibition, we used intracerebral drug microinfusion and chemogenetic DREADD (Designer Receptors Exclusively Activated by Designer Drugs; Roth, 2016) methods. First, we used mPFC microinfusion of the GABA-A receptor agonist muscimol or antagonist picrotoxin, respectively, which we had previously shown to induce mPFC inhibition/disinhibition, respectively, including reduced/enhanced neuronal burst firing (Pezze et al., 2014). Second, we expressed the inhibitory DREADD hM4Di in mPFC GABAergic neurons to inhibit these neurons (‘chemogenetic disinhibition’). Specifically, we Cre-dependently expressed hM4Di in the mPFC of Vgat-Cre rats. Because this approach is novel, we confirmed expression and functionality of the inhibitory DREADD using histological and in vivo electrophysiological methods before examining how such chemogenetic mPFC disinhibition affected reversal learning. To complement whole-session performance measures, including responses to criterion (RTCs) and errors, we used Bayesian trial-by-trial analysis of behavioural strategies (including win-stay and lose-shift) underlying reversal performance (Maggi et al., 2024). Because early reversals may require more mPFC-dependent attention and cognitive control (to overcome pre-existing response biases) than serial reversals, we predicted that mPFC functional inhibition may impair early reversals more than serial reversal learning. Additionally, we predicted that mPFC disinhibition may disrupt reversal performance, regardless of whether the mPFC is required, because disinhibition may disrupt processing in mPFC projection sites.

## MATERIALS AND METHODS

### Rats

In total, we used 101 young adult male rats all supplied by Envigo (UK). This included: two cohorts of Lister hooded rats for the mPFC microinfusion studies; one cohort of Lister hooded rats for a control study on the effect of the DREADD ligand clozapine-N-oxide (CNO) on reversal learning; and two cohorts of Vgat-Cre rats, for the chemogenetic studies (**Table 1**). The Vgat-Cre rats (homozygous LE-VGATem1(IRES-Cre)Sage; https://www.inotiv.com/research-model/hsdsage-le-vgatem1-ires-cre-sage) are Long-Evans hooded rats with a knock-in of Cre-recombinase under control of the endogenous VGAT (vesicular GABA transporter) promoter, to enable Cre expression in GABAergic neurons. Listerhooded rats weighed between 290-340 g (8-9 weeks), and Vgat-Cre rats weighed 300-460 g (8-11 weeks old) at surgery (cannula implantation or viral injection, respectively). Seven out of 64 Lister hooded rats used for the microinfusion studies had to be excluded: four rats required euthanasia due to complications with the chronic cannula implant; one rat did not regain consciousness after surgery; two rats were excluded due to an apparatus malfunction during operant testing. Two of 21 Vgat-Cre rats had to be excluded due to convulsive seizures, which were observed 35 days after viral vector injection and before any CNO injections, and two additional Vgat-Cre rats were excluded from the electrophysiological characterization involving the injection of 6.0 mg/kg CNO-2HCl, because the electrodes were located too anterior and there was insufficient DREADD expression around the electrode tips; brain sections of these two rats, which showed strong mPFC DREADD expression, could still be included in the histological analyses. Finally, one Vgat-Cre rat in cohort #4 (**Table 1**) showed repeated, but infrequent, seizures, and, therefore, did not require euthanasia or exclusion from testing and analysis. For sample size justifications, please see relevant sections below outlining the different experiments.

**Table 1.**
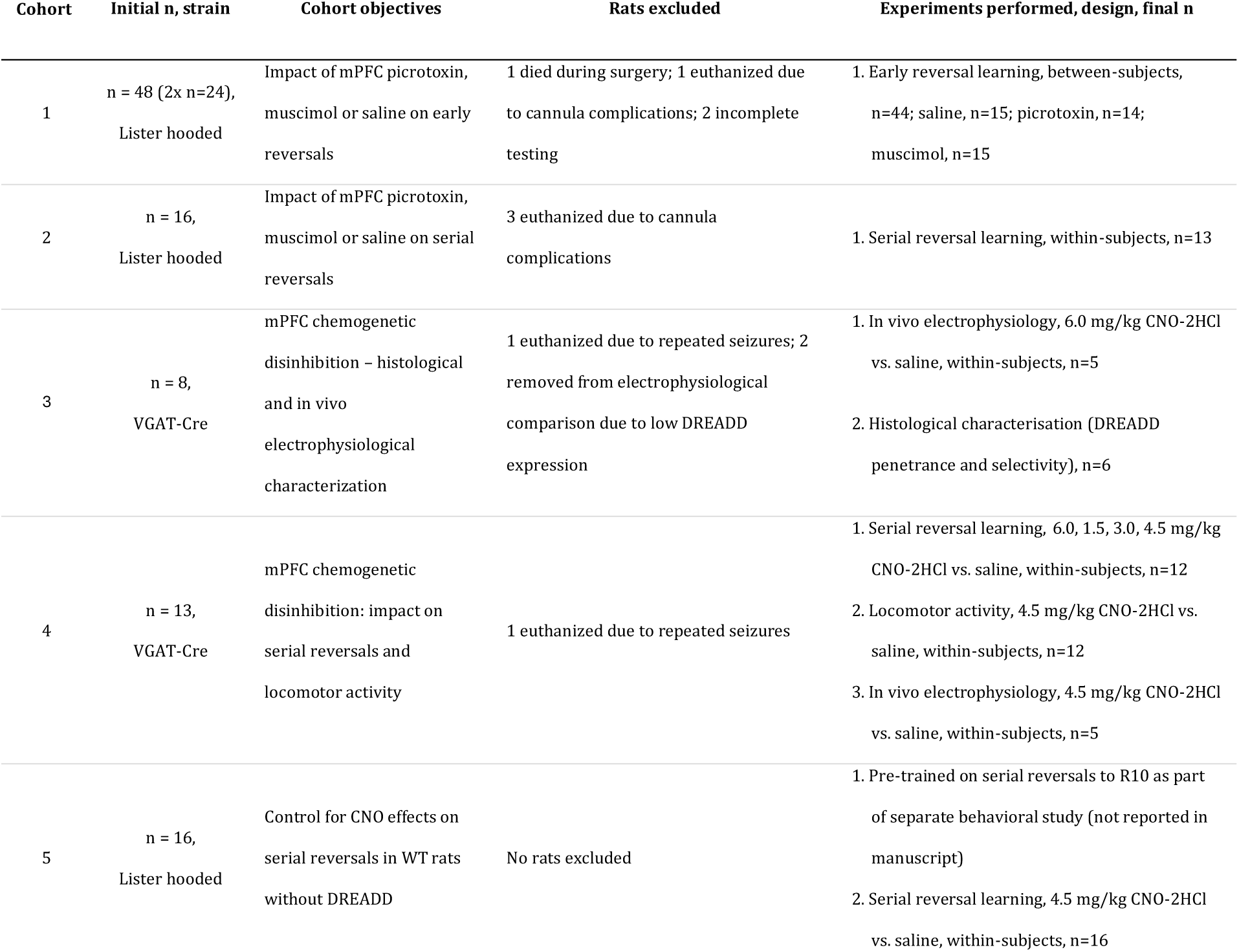
Rat cohorts used in the present study. Rats are divided into five cohorts by key experimental purpose. Initial rat numbers (n), exclusions and resulting final n after exclusions are shown. Experiments in cohort #1 were run in two batches of n=24 rats (total n=48). Note, one additional Vgat-Cre rat (cohort #4) showed repeated but infrequent seizures which did not require euthanasia, and, therefore, was not excluded.

| Cohort | Initial n, strain | Cohort objectives | Rats excluded | Experiments performed, design, final n |
| --- | --- | --- | --- | --- |
| 1 | n = 48 (2x n=24),<br>Lister hooded | Impact of mPFC picrotoxin,<br>muscimol or saline on early<br>reversals | 1 died during surgery; 1 euthanized due<br>to cannula complications; 2 incomplete<br>testing | 1. Early reversal learning, between-subjects,<br>n=44; saline, n=15; picrotoxin, n=14;<br>muscimol, n=15 |
| 2 | n = 16,<br>Lister hooded | Impact of mPFC picrotoxin,<br>muscimol or saline on serial<br>reversals | 3 euthanized due to cannula<br>complications | 1. Serial reversal learning, within-subjects, n=13 |
| 3 | n = 8,<br>VGAT-Cre | mPFC chemogenetic<br>disinhibition – histological<br>and in vivo<br>electrophysiological<br>characterization | 1 euthanized due to repeated seizures; 2<br>removed from electrophysiological<br>comparison due to low DREADD<br>expression | 1. In vivo electrophysiology, 6.0 mg/kg CNO-2HCl<br>vs. saline, within-subjects, n=5<br>2. Histological characterisation (DREADD<br>penetrance and selectivity), n=6 |
| 4 | n = 13,<br>VGAT-Cre | mPFC chemogenetic<br>disinhibition: impact on<br>serial reversals and<br>locomotor activity | 1 euthanized due to repeated seizures | 1. Serial reversal learning, 6.0, 1.5, 3.0, 4.5 mg/kg<br>CNO-2HCl vs. saline, within-subjects, n=12<br>2. Locomotor activity, 4.5 mg/kg CNO-2HCl vs.<br>saline, within-subjects, n=12<br>3. In vivo electrophysiology, 4.5 mg/kg CNO-2HCl<br>vs. saline, within-subjects, n=5 |
| 5 | n = 16,<br>Lister hooded | Control for CNO effects on<br>serial reversals in WT rats<br>without DREADD | No rats excluded | 1. Pre-trained on serial reversals to R10 as part<br>of separate behavioral study (not reported in<br>manuscript)<br>2. Serial reversal learning, 4.5 mg/kg CNO-2HCl<br>vs. saline, within-subjects, n=16 |

Rats were housed in groups of three or four, in individually ventilated ‘Double Decker’ cages (462 mm x 403 mm x 404 mm; Tecniplast, UK) with temperature and humidity control (21±1.5 °C, 50±8%) and 12-h light-dark cycle (lights on at 0700). All experimental procedures were carried out during the light phase. Rats had free access to water throughout the study. Access to food (Teklad Global 18% Protein Rodent Diet 2018C; Envigo, UK) was *ad libitum* until the start of food restriction one day prior to the start of operant pretraining. During food restriction, rats received daily food rations of 15-21 g per rat and were weighed daily to maintain body weights at 85-90% of free-feeding weights, as projected by a pre-established weight growth curve. On pretraining and test days, rats were weighed before the day’s operant task session and received their daily food ration after completing the session (in addition to the food reward received during the session). All in vivo procedures were conducted in the Biological Support Unit (BSU) at the University of Nottingham in accordance with the United Kingdom (UK) Animals (Scientific Procedures) Act 1986, approved by the University of Nottingham’s Animal Welfare and Ethical Review Body and run under the authority of Home Office project license PP1257468. Methods and results are reported in line with the ARRIVE guidelines (Lilley et al., 2020).

### mPFC functional inhibition and disinhibition by drug microinfusions

#### Guide cannula implantation into mPFC for microinfusions

Guide cannulae were implanted into the mPFC using the same coordinates and similar methods as in our previous studies (Pezze et al., 2014). Rats were anesthetized with isoflurane (induction: 3%; maintenance: 1-3%) delivered in medical oxygen (1 L/min). Following induction of anesthesia, rats’ scalps were shaved, and all rats received subcutaneous (s.c.) injections of analgesic (Rimadyl 50 mg/mL; Zoetis, UK, diluted 1:9 with sterile saline, 0.9%, and injected at 0.1 mL/100 g) and antibiotic (0.02 mL/100 g Synulox containing 14% Amoxicillin; Zoetis, UK). Rats were transferred to the stereotaxic frame where they were secured in the horizontal-skull position with ear bars coated with local anesthetic cream (EMLA 5%, containing 2.5% lidocaine and 2.5% prilocaine; AstraZeneca, UK). Eye gel (Lubrithal, Dechra, UK) was applied to the eyes to prevent drying out during surgery, and the shaved scalp was disinfected with alcoholic skin wipes (2% chlorhexidine, 70% alcohol; Clinell, UK). Throughout the surgery, body temperature was maintained at 37 °C via a homeothermic heating pad controlled by an external temperature probe placed under the rat. A small anterior-posterior incision was made into the scalp to expose the skull, and bregma was located. Two small holes were drilled through the skull at the following coordinates: +3 mm anterior and ±0.6 mm lateral from bregma. Bilateral infusion guide cannulae (“mouse” model C235GS-5-1.2; Plastics One, Bilaney Consultants, UK) were used, consisting of a 5-mm plastic pedestal that held two 26-gauge metal tubes, 1.2 mm apart and projecting 4.5 mm from the pedestal. Using a cannula holder attached to the stereotaxic frame, the guide cannulae were lowered to 3.5 mm ventral from the skull surface and secured to the skull with dental acrylic (Kemdent, UK) and stainless-steel screws (1.2 mm x 3 mm; MDK Fasteners, UK). Non-protruding double stylets (33 gauge; Plastics One, Bilaney Consultants, UK) were inserted into the guide cannulae and a dust cap was secured on top. At the end of surgery, rats were injected with 1 mL of saline (s.c.) to minimize the risk of dehydration. Following surgery, rats continued to receive daily prophylactic antibiotic injections (0.02 mL/100 g, s.c., Synulox containing 14% Amoxicillin; Zoetis, UK) for the duration of the study. Rats were allowed to recover for at least 7 days before the start of food restriction and pretraining on the operant task.

#### mPFC drug microinfusions

Muscimol and picrotoxin doses and infusion parameters were based on our previous studies, which showed that mPFC infusions of these drugs cause neural changes consistent with neural inhibition or disinhibition (including reduced or enhanced neuronal burst firing), as well as marked attentional deficits on the 5-choice serial reaction time task (Pezze et al., 2014). Rats were gently restrained to insert 33-gauge injectors (Plastics One, Bilaney Consultants, UK) into the previously implanted guide cannulae. The injectors protruded −0.5 mm below the guides into the mPFC, resulting in the following target coordinates for the infusions: +3 mm anterior and ±0.6 mm lateral from bregma and −4 mm ventral from skull. The injector ends were connected via polyethylene tubing to two 5-μl syringes (SGE, World Precision Instruments, UK) secured on a micro-infusion pump (sp200IZ, World Precision Instruments, UK). Prior to infusions, the tubing and syringe were backfilled with distilled water, and an air bubble was included before any drug solution was pulled up. A volume of 0.5 µL/side of either 0.9% sterile saline (vehicle), GABA-A receptor antagonist picrotoxin (300 ng/0.5 µL/side, C30H34O13, Sigma-Aldrich, UK) in sterile saline, or agonist muscimol (62.5 ng/0.5 µL/side, C4H6N2O2, Sigma-Aldrich, UK) in sterile saline was administered over the span of 1 min. Movement of the air bubble within the tubing was monitored to ensure solutions had been successfully injected into the brain. Injectors remained in place for an additional 1 min to allow for absorption of the infusion bolus by the surrounding brain tissue. The injectors were then removed, and the stylets replaced. Testing began 10 min after the infusion was complete. Each rat received six microinfusions into the mPFC. The number of microinfusions per rat was limited to six to reduce the risk that reactive gliosis caused by repeated microinfusions may reduce effectiveness of the infused drug (Cunningham et al., 2008). After each infusion session, tubing and microinjectors were flushed with distilled water, and cleaned with >98% ethanol.

#### Verification of cannula placements

At the end of all testing, rats were overdosed with sodium pentobarbitone (1–2 mL Euthatal; sodium pentobarbitone, 200 mg/mL; Genus Express, UK) and transcardially perfused with PBS followed by 4% paraformaldehyde (PFA) solution in PBS. Following extraction, brains were post-fixed in 4% PFA and cut into 70-µm coronal sections using a vibratome (Leica, UK). Sections were then mounted on slides and viewed under a light microscope, where cannula placements were verified and mapped onto coronal sections of a rat brain atlas (Paxinos and Watson, 1998). A subset of sections was cresyl-violet stained and coverslipped (DPX mountant, Sigma-Aldrich, UK) to take photographs for presentation purposes.

### Chemogenetic mPFC disinhibition

#### Viral vector injections into mPFC

We used a double-floxed Gi-coupled hM4Di DREADD fused with mCherry reporter under the control of human synapsin promoter and packed in an Adeno Associated Virus (AAV) vector (pssAAV-8-hSyn1-dlox-hM4D(Gi)_mCherry(rev)-dlox-WPRE-hGHp(A), https://www.addgene.org/44362/). For injections, we used a solution of the DREADD-containing viral vector, serotype 8 (CNS tropism), in 1x PBS, pH 7.4 (including 1 mM MgCl_2_ and 2.5 mM KCl), with a physical titre of 6.3×10^12^ vg/mL (v84-8, Viral Vector Facility, University of Zürich, Switzerland). The solution was aliquoted into 12.5 µL portions into low protein binding PCR tubes and stored at −80 °C until use.

We used similar stereotaxic procedures and perioperative care as for the guide cannula implants described above; because a chronic implant was not required, rats did not receive any antibiotic treatment. Coordinates and injection parameters were based on pilot studies (chapter 4 in Renström, 2024). Skull trepanations were drilled above the mPFC at +2.8 mm anterior and ±0.6 mm lateral from bregma. Double injectors (C235I-SPC, Plastics One, Bilaney Consultants, UK) glued into double infusion guide cannulae (“mouse” model C235GS-5-1.2; Plastics One, Bilaney Consultants, UK, with 4.0 mm projection) were used to inject the vector. The injectors were held by a cannula holder attached to the stereotaxic frame. The ends of the injectors were attached to two 5 μL syringes (SGE, Australia) via PE50 tubing (Plastics One, Bilaney Consultants, UK), and syringes were secured on a micro-infusion pump (SP200IZ syringe pump, World Precision Instruments, UK). Prior to injections, the tubing and syringe were backfilled with distilled water, and a small air bubble was pulled into the tubing before any viral solution was pulled up. The injector tips were slowly lowered through the trepanations to −4.0 mm from the skull surface and then pulled back to the target dorsoventral coordinate of −3.8 mm, to create a small ‘pocket’ for the injection bolus. Rats received 1.0 µL of the AAV-solution bilaterally at a rate of 0.1 µL/min. Following injections, the injectors were left in place for 10 min to allow for absorption of the injection bolus. After the injectors had been withdrawn, the incision was sutured up. Rats were held for 28 days before any further procedures were conducted, to allow for strong DREADD expression (Smith et al., 2016).

#### CNO injections

We used systemic injection of CNO for DREADD activation. Previous studies have typically used 5-10 mg/kg of freebase CNO to activate DREADDs (Smith et al., 2016). For the DREADD activation in the current experiment we used CNO-2HCl (synthesised in-house, Boehringer Ingelheim, Germany), a water-soluble CNO salt. Sterile saline (0.9%) was used as vehicle and all systemic injections were intraperitoneal (i.p.), in a volume of 1 mL/kg. We started with an initial dose of 6 mg/kg of CNO-2HCl, which corresponds to 5 mg/kg of freebase CNO(molecular weight of CNO-2HCl compared to CNO:415 g/mol vs. 342 g/mol, a factor of 1.2). At 6.0 mg/kg, CNO-2HCl produced marked in vivo electrophysiological changes under anesthesia, in the mPFC of Vgat-Cre rats expressing the hM4Di DREADD in mPFC GABA neurons (**Fig. S1**). However, testing the effects of 6 mg/kg CNO-2HCl on reversal learning performance in Vgat-Cre rats expressing hM4Di in mPFC GABA neurons revealed that this dose had substantial side effects, with rats not pressing the operant levers. We, therefore, tested lower doses of 1.5, 3.0 and 4.5 mg/kg CNO-2HCl. Within the Results section, we focus on the results obtained with the 4.5 mg/kg dose. See below for additional details on the dose selection (Reversal learning experiments, Impact of mPFC muscimol and picrotoxin microinfusion and of mPFC chemogenetic disinhibition on serial reversal learning performance).

#### Verification of DREADD expression

At the end of all testing, rats were overdosed with sodium pentobarbitone (1–2 mL Euthatal; sodium pentobarbitone, 200 mg/mL; Genus Express, UK) and transcardially perfused with PBS followed by 4% paraformaldehyde (PFA) solution in PBS. Following extraction, brains were post-fixed in 4% PFA and cut into 50-µm coronal sections using a vibratome (Leica, UK). Sections were then mounted on slides (DPX mountant, Sigma-Aldrich, UK) and scanned under a fluorescence microscope (Zeiss AxioScan Z1 scanner; Carl Zeiss, Germany), to map expression of the mCherry reporter that was included with the DREADD. DREADD spread was visualised using Affinity Designer (Version 1.10.4, Serif, UK). Scans showing mCherry signal were superimposed onto a digital template of the Paxinos and Watson (1998) rat atlas. Due to variability in brain shape and size, scans were manipulated to best fit the template at the approximate distance from bregma. mCherry signal was traced by hand, and opacity of the regions reduced to 25% such that overlapping regions appeared darker.

### Reversal learning experiments

#### Food reinforced two-lever operant reversal task

The pretraining and testing protocols were adapted from our previous studies (Gonçalves et al., 2023) and based on original protocols by Brady and Floresco (2015).

##### Apparatus

All reversal testing was conducted in eight individual operant boxes (30.5 cm x 24.1 cm x 21.0 cm; Med-Associates, St. Albans, VT, USA), equipped with a house light (40 lux), an extractor fan, two retractable response levers either side of a circular dish, into which sucrose reward pellets (5TUL-45mg, Testdiet, OCB Systems Ltd, UK) were dispensed, as well as two LED cue lights (40 lux each), one above each lever. The LED cue lights above the levers were illuminated pseudo-randomly throughout all sessions but were not relevant to the correct lever choice (which was always determined by lever position, left or right). Each rat was assigned to an operant chamber, where it underwent all operant pretraining and test sessions. Chambers were cleaned with 20% ethanol between different rats. The stimuli presented, lever operation and data collection were controlled via an interface with the computer and using custom software (MED-PC software, Med-Associates, St. Albans, VT, USA).

##### Pretraining

One day before pretraining, rats were habituated to the apparatus by placing them in the operant box for 15 min, with exterior doors open and no levers extended. Pretraining started with several days of single lever-press training sessions (one session per day), during which one of the two levers was extended for a fixed 30-min period, with each lever press rewarded by one sucrose pellet (5TUL-45mg, Testdiet, OCB Systems Ltd, UK). The initial location (left or right) of the lever was counterbalanced across rats. Lever location was switched once at least 50 responses were made in one session. At the beginning of the first lever-press session, two pellets were placed in the dish and crushed pellets on top of the extended lever to promote rats’ engagement with the apparatus. This was repeated on day two if rats did not readily press the lever on the previous day (<10 presses in the first 10 min). During these sessions, rats could press levers without restriction, allowing for unlimited rewards. This stage was completed after the criterion of 50 responses was met for both levers.

Lever-press training was followed by a minimum of 5 days of 90-trial retractable-lever training sessions (one session per day), where rats had to press the extended lever within the 10-s response window, after which the lever would be retracted. Both levers were presented in a pseudo-random order, but the same lever would not be extended more than two times in a row. Each lever press was rewarded by one sucrose pellet. At the end of this stage, rats were expected to make 5 or fewer omissions during a 90-trial session. 51/64 rats achieved this criterion by the last day. All rats had achieved several consecutive days of <5 omissions prior to the last day, with all rats making an average of 3.67±0.58 (x̄SEM) omissions by the final day, and thus all rats progressed to the next stage.

On the final day of pretraining, immediately after the last 90-trial session, the side preference of each rat was determined via 7 trials consisting of several sub-trials. At the start of each main trial, both levers were extended into the chamber, and the initial press was rewarded, regardless of location. Subsequent sub-trials only rewarded the first response on the opposite lever. Once the rat had pressed the opposite lever, the next main trial began. Therefore, each of the 7 trials included one rewarded press on each lever. The rat’s side preference was defined as the side of the lever, which the rat chose to start on at least 4 of the 7 main trials.

##### Spatial discrimination and reversal testing

Testing began with a spatial discrimination (SD) task, where rats were rewarded to press the lever opposite to their side preference established on the previous day. Trials began every 20 s, with a 10-s response window. On each trial, both levers were presented, but only the one opposite to the rats’ side bias was rewarded. Each correct response was rewarded with one sucrose pellet. Sessions were terminated once rats had reached a criterion of 10 consecutive correct responses. If the criterion was not reached within 200 trials, rats were retested with the same lever the next day. In the present study, all rats reached criterion within a maximum of two SD sessions.

The day after completion of the SD stage, reversal testing began. Each reversal stage started with 20 reminder trials, where the same lever response was rewarded as on the previous day. The purpose of reminder trials was to assess if retrieval/expression of the rule learnt on the previous day was affected by the prefrontal manipulations and, thereby, to aid interpretation of any effects of these manipulations on reversal learning (see Brady and Floresco, 2015). The reversal occurred after trial 20, when the correct and incorrect levers were reversed, such that the lever opposite to the one rewarded on the previous day and during the reminder trials was now the correct lever and rewarded. A reversal session ended when rats reached a criterion of 10 consecutive correct responses or after a maximum of 200 reversal trials. The protocol for unsuccessful sessions, where the success criterion was not attained, differed between the experiments examining early reversals and serial reversals and is described in the following two sections.

#### Impact of mPFC muscimol and picrotoxin microinfusion on early reversal learning performance

We used a between-subjects design to examine the impact of saline, picrotoxin and muscimol infusion on rats’ performance during the SD stage and during the first five reversals (R1 to R5). Rats (**Table 1**, Cohort #1) were allocated to one of the three infusion groups - saline, picrotoxin or muscimol - via a randomized block design, with at least one rat in each cage of four being allocated to each infusion group. Our target sample size was *n*=16 rats per group to give a power of 80% for pairwise comparisons (two-tailed independent t-tests, *p*<0.05), to detect differences between infusion groups that correspond to an effect size of Cohen‘s *d* of around 1 (G*Power, Faul et al., 2009). Rats received mPFC microinfusions 10 min before the SD and each reversal session, based on the onset of behavioral and electrophysiological effects of mPFC muscimol and picrotoxin infusion in our previous studies (Pezze et al., 2014). If rats did not reach the criterion of 10 consecutive correct trials within a maximum of 200 trials, they continued to be tested at the same stage (i.e., with the same lever rewarded), on the following day, 10 min after receiving their mPFC infusion. Because we limited the number of infusions per rat to six, rats that required more than one daily 200-trial session to complete a reversal stage, would complete less than five reversal stages. Therefore, statistical analysis was restricted to SD and R1-3, as these were the stages completed by all rats, with only 8/15 rats in the picrotoxin group successfully completing all 5 reversals.

#### Impact of mPFC muscimol and picrotoxin microinfusion and of mPFC chemogenetic disinhibition on serial reversal learning performance

Where we examined the impact of mPFC manipulations (drug microinfusions or chemogenetic; **Table 1**, Cohort #2 and #3) on serial reversals, rats first underwent the SD stage and four reversals (R1 to R4), to establish relatively stable reversal performance, based on previous pilot work (chapter 2 in Renström, 2024), before the impact of mPFC manipulations was tested within-subjects across subsequent reversals stages. Our target sample size was *n*=12 rats to give a power of >80% for pairwise comparisons (two-tailed paired t-tests, *p*<0.05) to detect differences between manipulation conditions that correspond to an effect size of Cohen‘s d of around 1. Within each session, rats were allowed maximally 200 reversal trials to reach criterion, and all rats reached criterion within this number of trials. To reduce the risk of carry-over effects, whereby any impairment caused by an mPFC manipulation during one reversal may affect performance on the next reversal and, thereby, confound the within-subjects design, we interspersed ‘retention days’ between reversals. On retention days, rats were tested without any drug microinfusion or CNO-2HCl injection on the same reward contingency as on the preceding infusion/injection day (i.e., during the last reversal) until they reached the criterion of 10 consecutive correct responses or for 200 trials (all rats reached criterion within less than 200 trials on retention days). Retention days reinforced the lever press rule from the last reversal before testing rats on the next reversal, to ensure that all rats started each new reversal with the old rule from the previous stage well established.

The impact of mPFC microinfusions of muscimol and picrotoxin was examined across R5 to R10. Each rat received two series of the three different mPFC infusions (saline, picrotoxin, and muscimol), with infusion series 1 consisting of infusions applied 10 min before R5-7 and infusion series 2 consisting of infusions applied 10 min before R8-10.

The impact of chemogenetic mPFC disinhibition by CNO-2HCl injection of Vgat-Cre rats expressing hM4Di in mPFC GABA neurons was also examined from R5 onwards, with testing order of the different injections counterbalanced using a Latin-square design (**Table 1**, Cohort #4). We started comparing the impact of 6 mg/kg of CNO-2HCl and saline injections across R5 and R6. Chemogenetic mPFC with 6 mg/kg of CNO-2HCl had substantial gross side effects: rats were very sensitive to handling and hardly pressed the levers during the 90-min test session. We, therefore, tested lower doses of CNO-2HCl. A comparison of the impact of 1.5 and 3.0 mg/kg, and saline, across R7 and R9, did not reveal any gross side effects, but also no significant impairments in whole-session measures of reversal performance (including RTCs and errors), although a Bayesian trial-by-trial analysis revealed a dose-dependent reduction in win-stay probability around reversal (**Fig. S2**). We, therefore, increased the dose to 4.5 mg/kg and compared this with saline at R10 and R11, which induced marked reversal impairments without inducing non-specific side effects. Therefore, this report will focus on the 4.5 mg/kg dose in the *Results* section.

Finally, to verify that CNO-2HCl did not affect serial reversal learning independent of DREADD activation, e.g. due to known back metabolism into clozapine (Jendryka et al., 2019), we compared the impact of systemic saline vehicle and 4.5 mg/kg CNO-2HCl injections on serial reversals in wild-type Lister hooded rats (**Table 1**, Cohort #5). These rats had undergone serial-reversal testing up to R10, for the purpose of another behavioral study. Then, at R11 and R12, we compared behavior following i.p. injection of 4.5 mg/kg CNO-2HCl or saline in a cross-over within-subjects design.

#### Blinding

In all reversal learning experiments the main experimenter (JR) was blinded with respect to the manipulation groups and conditions. Blinding was only broken after data analysis (although the identity of the picrotoxin group/condition and the 6 mg/kg CNO-2HCl condition was evident from the increased omissions).

#### Performance measures

##### Whole-session measures

The main measure of operant task performance was RTC, i.e., the number of total trials a rat required to achieve the success criterion of 10 consecutive correct responses, excluding omitted trials when the rat did not respond within the 10-s response window. For the reminder trials, which continued even if rats made 10 consecutive correct responses, ‘percentage correct’ was used, which refers to the percentage of correct reminder-trial responses divided by the total number of reminder-trial responses, excluding omissions. For the analysis of errors during reversal sessions, previous studies used different approaches to differentiate between ‘perseverative errors’, which reflect difficulty in abandoning the previously learned rule, and ‘regressive errors’, which occur when the old rule has been abandoned but performance slips back due to errors in performing the new rule (Brady & Floresco, 2015; Dalton et al., 2016; Deuel et al., 1971; Dhawan et al., 2019; Floresco et al., 2006). We compared several different approaches, and all yielded similar conclusions. Here, we only report the outcome of the approach outlined in Brady and Floresco (2015). We counted errors as perseverative errors until a threshold of less than 10 total errors in a block of 16 responses was reached. After this, all errors were counted as regressive. Additionally, we analyzed omissions (i.e., trials when rats did not press a lever within the 10-s response window), as well as response latencies for correct and incorrect responses. Performance measures during SD and reversal trials were analyzed separately from the performance measures during the reminder trials at the beginning of each reversal stage.

##### Trial-by-trial Bayesian strategy analysis

Complementing the classical performance measures outlined above, we also used a trial-by-trial Bayesian strategy analysis (Maggi et al., 2024). This analysis estimates the probability of a rat applying a particular response strategy on any given trial based on evidence collected up to this trial. Each strategy (e.g., lose-shift, win-stay) was estimated independently from each other based on a set of input parameters. These parameters included the choice (i.e., right or left lever) and the outcomes (i.e., correct/rewarded or incorrect/unrewarded) of the current and preceding responses. For example, a ‘lose-shift’ strategy is defined as an incorrect/unrewarded response (‘lose’), followed by a response ‘shift’ to the opposite lever on the following trial, whereas a ‘win-stay’ strategy is defined as a correct/rewarded response (‘win’), followed by re-selection of the same lever (‘stay’) on the following trial. Estimating the lose-shift/win-stay strategy probabilities, therefore, requires information about the choice and outcome on trial t and the choice on trial t+1. Briefly, the Bayesian strategy analysis uses the Bayes rule to estimate the probability of a given strategy *i* being applied by the subject based on the history of choices up to the current trial *t,* P(strategy_i_(t)|choices(1:t)). This *posterior* probability is updated based on the *likelihood* (how well the choices match the strategy) and the *prior* (initial probability estimate of using that strategy). The initial estimation of the prior at trial 0 was based on the uniform distribution. Maggi et al. (2024) showed that the choice of prior (whether based on the uniform or Jeffrey’s distribution) has minimal impact on the estimated strategy probabilities after just a few trials because the actual data quickly dominates over the initial prior assumptions. At subsequent estimations, probabilities at trial *t-1* were used as priors for posterior estimations at trial *t*. The model also included a forgetting rate parameter, which was not modified from our previous study and progressively reduced the influence of past evidence while giving more weight to recent choices in the probability calculations; moreover, if no evidence was available for or against a given strategy on a particular trial, the strategy’s probability remained the same as on the previous trial (Maggi et al., 2024). To establish a robust estimation of strategy probabilities in the study of early reversals, we used the SD stage to accumulate evidence for each strategy before statistically comparing the impact of mPFC manipulations on probabilities at all stages from R1 onwards. In the study of serial reversals, evidence for each strategy was accumulated across the SD stage and R1-R4 before comparing the impact of mPFC manipulations on probabilities from R5 onwards.

We investigated strategies around reversal (i.e., responses immediately preceding and following the rule reversal) and preceding learning (i.e., responses preceding the 10 consecutive correct responses that we defined as criterion of successful learning and that ended the test sessions). These time points were of interest, as response strategies around rule reversal reveal how the rats respond to the rule change, whereas strategies preceding learning reveal how rats ultimately learn the new rule. The number of responses included in our Bayesian strategy analysis was limited by the number of responses that were completed by all rats across all sessions. For the mPFC microinfusion studies, strategy analysis prior to rule reversal was limited to 6 reminder-trial responses. We kept this consistent for the analysis of the chemogenetic study, although in the chemogenetic study more reminder trials would have been available for strategy analysis because chemogenetic disinhibition caused less omissions. For both microinfusion studies, 16 responses following reversal were included (i.e., overall 22 responses around rule reversal, 6 before and 16 after reversal), but because some rats in the chemogenetic study reached criterion in only 10 responses, we could only include 10 responses after reversal in that analysis (i.e., overall 16 responses around rule reversal, 6 before and 10 after reversal).

As the last 11 responses of every session were identical (one incorrect response preceding the success criterion of ten correct responses), our analysis of strategies preceding learning focused on the last 5 responses before the last incorrect response in that session. In the chemogenetic study, due to several rats achieving criterion within 10 responses after reversal, we could not analyse strategies around learning, because in these cases the defined point of learning (first of the 10 successive correct response to reach criterion) was identical with the point of rule reversal and, therefore, there were no responses prior to learning after rule reversal.

We examined the following strategies:

1. Strategies that reflect if rats follow the task rule - ‘Go-previous’ strategy around reversal, and its complementary strategy ‘go-new’ around learning: The go-previous strategy involves the rat following the spatial rule that was correct prior to rule change (i.e., go-right or go-left), whereas the go-new strategy involves the rat applying the correct choice strategy after the reversal. Go-previous and go-new strategies are mutually exclusive, i.e. their probabilities add up to P=1, whereby as one increases the other one decreases, and vice versa. We focused on go-previous around reversal and go-new around learning, because these provide information about how flexibly rats moved away from the previously learned rule and how quickly they learned the new rule, respectively.
2. Other explorative and exploitative strategies: Win-stay and lose-shift strategies are both task-appropriate strategies that would support correct responding, because the rat continues to press the correct lever or ‘shifts’ to the other lever after an incorrect choice, respectively. In addition, to capture alternative approaches that rats may use to adjust to the reversal, we also examined strategies that were task-inappropriate, such as cue-light-based strategies, and situational strategies that are neither directly appropriate nor inappropriate, such as a ‘sticky’ strategy, which involved ‘sticking’ with the same response regardless of outcome. We only report these probabilities if they varied substantially from chance (P=0.5). Finally, for all the strategies above, there is a strategy that is mutually exclusive (go-previous and go-new; win-stay and win-shift; lose-shift and lose-stay; sticky and alternating). Because the probability of one of these two mutually exclusive strategies is implied by the probability of the other, we only analyzed the probabilities of the dominant strategy (i.e., P>0.5) out of these pairs of mutually exclusive strategies.

### Histological, in vivo electrophysiological and behavioural characterization of DREADD expression and functionality in the mPFC

#### Histological analysis of the spread, penetrance and specificity of DREADD expression

A cohort of Vgat-Cre rats (n=5; **Table 1**, Cohort #3) was used to confirm suitable DREADD expression across the mPFC. More specifically, we examined three aspects of DREADD expression: 1) the extent of DREADD spread across the mPFC and to adjacent non-mPFC regions; 2) DREADD penetrance (i.e., which proportion of GABAergic neurons expressed the DREADD); 3) cell-type specificity of the DREADD (i.e., if DREADD expression was largely limited to GABAergic neurons). For this purpose, rats were overdosed with sodium pentobarbitone (1–2 mL Euthatal; sodium pentobarbitone, 200 mg/mL; Genus Express, UK), 28 days following mPFC viral vector injection, and transcardially perfused with 160 mL of freshly prepared PBS followed by 160 mL PFA (4%). After extraction, brains were post-fixed in 4% PFA overnight at +4°C, transferred into sucrose solution (30%) at +4°C and shipped to facilities at Boehringer Ingelheim, Biberach, Germany, where they were cut into 50-µm coronal sections on a sliding microtome (Microm HM450, Thermo Fisher Scientific, Germany) with integrated freezer platform (BFS-3MP, Physitemp, Germany), and immediately placed into wells (24-well, Nuclon Delta Surface; Thermo Fisher Scientific, Germany) filled with 2 mL PBS and 0.01% thimerosal. Brain sections were stored at +4°C until further processing. For histological verification of DREADD expression spread (1), due to the endogenous fluorescence of the mCherry reporter no additional staining was required, so slices were mounted using 1-2 drops of ProLong Diamond Antifade Mountant containing DAPI (Invitrogen, USA) and coverslipped, before being scanned on a florescence microscope (Zeiss AxioScan Z1 scanner; Carl Zeiss, Germany) using a 20x objective (0.22 µm/px) controlled by ZEN software (Carl Zeiss) equipped with a Zeiss Colibri 7 LED light source (Carl Zeiss). To quantify DREADD penetrance (2) and cell-type specificity (3) we used immunohistochemical staining procedures. Blocking solution (1% Bovine serum albumin (BSA), 0.3% Triton-X 100, 10% Normal goat serum (NGS; diluted in PBS)) was applied to the slices and incubated at room temperature for 2 h. Then, primary antibodies diluted in an antibody solution (1% BSA, 0.3% Triton-X 100, 1% NGS) were added and the slices were incubated at +4 °C overnight. Primary antibodies included anti-glutamate decarboxylase 67 (GAD67; rabbit, Millipore, Cat#MAB5406) labelling GAD67, a key synthesis protein of GABA, anti-calmodulin dependent protein kinase II alpha (CaMKIIa, rabbit, Novus Biologicals Cat#NBP2-46821) labelling glutamatergic neurons, anti-choline acetyltransferase (ChAT, rabbit, Synaptic Systems Cat#297013) labelling cholinergic neurons, and anti-red fluorescent protein (RFP, mouse, Synaptic System Cat#390011) used to label and thus amplify endogenous mCherry signal. Following washing (45 min), via a washing solution (PBS, 0.1% Triton-X 100), secondary antibodies conjugated to Alexa 488 and 568 (Thermo Fisher Scientific, Germany), were applied and slides were incubated at room temperature for 3 h in the dark. Following another wash (60 min), 1-2 drops of ProLong Diamond Antifade Mountant containing DAPI (Invitrogen, USA) were applied to each slide and slides were cover slipped. Slices stained immunohistochemically were scanned on the Opera Phenix high-content confocal microscope (PerkinElmer, USA) running Harmony high-content imaging software (PerkinElmer) using a 63x water immersion objective (0.19 μm/px). We used four slices per rat for GAD67 and one slice per rat for CaMKIIa, as the latter was predominantly a control staining to ensure no expression in excitatory cells.

DREADD spread was visualised as described above (Chemogenetic/DREADD methods, Verification of DREADD expression). For cell classification and quantification, scans of the immunohistochemically stained brain sections were exported into HALO (IndicaLabs, USA) running the Multiplex FISH module. Parameters for automated cell classification were determined by examination of unambiguously GAD67+, mCherry+, ChAT+, and CaMKIIa+ cells. Subsequently, each slice was examined for potential false positive cases of cell classification, and the inclusion threshold was adjusted to best account for unambiguously positive cells, whilst minimizing the inclusion of false positive cases. Finally, each slice was examined once more to confirm parameters were suitable before automated cell counting was started.

#### In vivo electrophysiology

We used in vivo electrophysiological recordings under anaesthesia to examine if chemogenetic mPFC disinhibition would cause numerical changes in LFP and multi-unit measures of mPFC neural activity, similar to those we had previously observed following mPFC picrotoxin (Pezze et al., 2014). First, electrophysiological recordings were conducted in the same Vgat-Cre rats (n=5) that we used for the initial histological characterization of DREADD expression (see above; **Table 1**, Cohort #3), using 6.0 mg/kg CNO-2HCl. Second, after our reversal study indicated that 6.0 mg/kg caused non-specific behavioral effects, whereas 4.5 mg/kg CNO-2HCl specifically disrupted serial reversal learning, we re-examined the effects of 4.5 mg/kg in a subset of rats that contributed to the reversal study after all behavioral testing was completed (n=5; **Table 1**, Cohort #4). In the Results section,we focus on the 4.5 mg/kg dose.

Rats were anesthetized with isoflurane (induction: 3%; maintenance: 1-3%) delivered in medical oxygen (1 L/min). After induction of anaesthesia, rats’ scalps were shaved. Rats were transferred to the stereotaxic frame where they were secured in the horizontal skull position, with ear bars coated with local anaesthetic cream (EMLA 5%, containing 2.5% lidocaine and 2.5% prilocaine; AstraZeneca, UK), eye gel (Lubrithal, Dechra, UK) was applied to the eyes to prevent drying out during the experiment, and the shaved scalp was disinfected with alcoholic skin wipes (2% clorhexedine, 70% alcohol; Clinell, UK). Throughout the experiment, body temperature was maintained at 37 °C via a homeothermic heating pad controlled by an external temperature probe placed under the rat. Following a scalp incision, the skull was removed over the target area above the right mPFC. The exposed dura was incised and removed via forceps. Throughout the duration of the recording session the cortex was kept moist with 0.9% saline.

A four-channel microwire array (four 50-µm Teflon-coated stainless-steel electrodes, spaced 0.25 mm apart and arranged in one row spanning ∼1 mm, with a stainless-steel ground wire; MicroProbes for Life Science, US) was then implanted into the right mPFC. The array was connected via a head stage to the recording system (see below, *Multi-unit and LFP recordings*). The array was fixed to the arm of the stereotaxic frame, such that the row of four microwires was arranged parallel to the midline of the brain, and was slowly lowered into the brain, with the centre of the array aimed at the following target coordinates: +3.2 mm anterior and +0.6 mm to the right from bregma and −4.0 mm ventral from the skull.

The electrode array was connected via a unity-gain multichannel head stage to a multichannel preamplifier (Plexon, US). The analogue signal was amplified (1000x) and filtered, via a band-pass filter, into multi-unit spikes (250 Hz to 8 kHz) and LFP signals (0.7 to 170 Hz). All recordings were made against ground, with the ground wire attached to the ear bar of the stereotaxic frame. The analogue signal was further amplified via a multichannel acquisition processor system (Plexon) (final gain up to 32,000). This system also provided additional filtering of multi-unit data (500 Hz to 5 kHz), digitalisation of spikes at 40 kHz, and LFP data at 1 kHz. Both, multi-unit and LFP data were viewed online with Real-Time Acquisition System Programs for Unit Timing in Neuroscience (RASPUTIN) software (Plexon). Using RASPUTIN, neural activity was recorded for 30 min for both baseline and post-saline injection, and 3 h post-CNO-2HCl injection. Multi-unit spikes were recorded when a predefined threshold of −240 µV was exceeded, and LFP data was recorded continuously.

Positioning of the electrodes at the target coordinates was followed by a stabilisation period of at least 30 min, during which anaesthesia was adjusted to a stable level (50±10 breaths per min, around 1.5-2% isoflurane). Following a 30-min recording of baseline neural activity without any injection, rats received an i.p. injection of saline (0.9%; 1 mL/kg), and neural activity was recorded for another 30-min period; then rats received an i.p. injection of CNO-2HCl and neural activity was recorded for 3 h following the injection. For the i.p. injections, the rat’s right hindlimb was gently raised to inject into the lower abdomen, the hindlimb was gently lowered again, and the start and end times of the injection were noted to accurately identify the start and end points of the pre- and post-injection periods.

Following the post-CNO-2HCl period during electrophysiological recordings, a current (1 mA, 10 s) was passed through the first and fourth microwire to create an electrolytic lesion at the tip of the electrode and mark its position. Following this marking, the electrode assembly was removed from the brain. Subsequently, pentobarbital was administered for a deeper anesthesia, rats were transcardially perfused as above, and brains were extracted and cut into 70 µm coronal sections, which were used to visualise spread of DREADD expression as described above and to map locations of marked electrode tips onto coronal sections of the rat brain atlas (Paxinos & Watson, 1998) using a light microscope (Leica DM 1000; Leica Microsystems, Germany).

Multi-unit and LFP data were analysed using Neuroexplorer (Version 3.2.2.6 Nex Technologies), as in our previous study (Pezze et al., 2014). Parameters of multi-unit activity (overall firing rate, number of bursts, percentage of spikes fired in bursts, mean within-burst firing rate, mean burst duration, inter-burst interval) and LFP power, as reflected by the area under the curve (AUC) of the power spectral density (PSD) function, were calculated in 5-min blocks across the whole recording session, including baseline, post-saline and post-CNO-2HCl periods. Prefrontal bursts were identified via the Poisson surprise method (Legendy & Salcman, 1985; Pezze et al., 2014), defining bursts as ‘improbable’ compared to the rest of the analysis window. LFP AUC of PSD was calculated by applying a Fast Fourier Transform analysis to the LFP signal from 0.7-170 Hz. All data was normalized to baseline by dividing values from each channel by the average value obtained from the same channel during the six 5-min baseline recording blocks. Normalized values were averaged across all 4 channels to obtain one single value of each parameter per 5-min block for each rat, resulting in 36 5-min blocks per rat (6 baseline, 6 post-saline and 24 post-CNO-2HCl).

#### Open-field measurement of locomotor activity

Using the same rats we used to examine the impact of mPFC chemogenetic disinhibition on serial reversals (**Table 1**, Cohort #4, n=12), we investigated the impact of mPFC chemogenetic disinhibition on locomotor activity in an open field, after all reversal testing had been completed. The open-field locomotor measurements were completed as described previously in our studies of mPFC disinhibition by picrotoxin (Pezze et al., 2014). In brief, locomotor measurements were conducted in 12 clear Perspex chambers (39.5 cm x 23.5 cm x 24.5 cm), surrounded by a two-level frame containing 12 photobeams (4 x 8 configuration; Photobeam Activity System, San Diego Instruments, US) and placed in a dimly illuminated room (50-70 lux). Two consecutive breaks of adjacent beams within the lower level of photobeams generated one locomotor count. Total locomotor counts were calculated for each 10-min block of testing. Sessions started automatically once rats were placed in the chambers and a first beam break was recorded. The effect of saline and 4.5 mg/kg CNO-2HCl injections on locomotor activity was compared in Vgat-Cre rats expressing the inhibitory DREADD in GABAergic neurons in the mPFC, using a within-subject design. Activity was measured across 5 successive days: Day 1 was a baseline/habituation day, on which activity was recorded for a 60-min period without any manipulations. Day 2 and 4 were ‘injection days’, beginning with 30 min of baseline recordings, after which rats received intra-peritoneal injections of either CNO-2HCl or saline. Following injections, rats were placed back in their chambers and activity was recorded for 60 min. Testing order of saline and CNO-2HCl injections was counterbalanced across rats. Days 3 and 5 were re-baseline days on which activity was recorded for a 60-min period, without any injection (as on day 1); re-baseline days served to detect any carry-over effects from the preceding injection days.

Locomotor counts were binned in 10-min blocks, giving six 10-min blocks for each baseline day, and nine 10-min blocks for each injection day (3 pre-injection and 6 post-injection blocks). These values were subsequently averaged across rats for each 10-min block.

### Statistical analysis

All statistical analyses were carried out in JASP (version 0.19.3), and all graphs were made in GraphPad prism (version 10.4.2). The accepted level of significance was p<0.05 for all analyses.

For the study examining the impact of mPFC muscimol or picrotoxin on early reversals, whole-session performance measures were analysed by ANOVA, using infusion group (saline, picrotoxin, muscimol) as a between-subjects factor and task stage (SD, R1-3) as a within-subjects factor. For the study examining the impact of mPFC muscimol or picrotoxin on serial reversals, whole-session performance measures were analysed by ANOVA using infusion condition (saline, picrotoxin, muscimol) and infusion series (1 or 2) as within-subject factors. For the studies examining the impact of chemogenetic mPFC disinhibition on serial reversals and locomotor activity, systemic injection (CNO-2HCl or saline) was used as a within-subjects factor, and 10-min block was used as an additional within-subjects factor for the locomotor data. For all reversal learning studies, trial-by-trial strategy probabilities were analysed by ANOVA, as outlined for the whole-session performance measures, and response number was used as an additional within-subjects factor (we refer to ‘responses’, instead of ‘trials’, due to the removal of omission trials in these analyses). Any significant interactions observed were further examined by simple-main effects analyses, and simple main effects were further analyzed by pairwise comparisons, using Fisher’s LSD test. Where a significant interaction was observed between multiple factors, the main effects of these factors were not reported, as these were obsolete given the interactions involving these factors. A statistical comparison of LFP and multi-unit measurements following CNO-2HCl or saline injection was not originally planned. However, to provide statistical support for the observed numerical changes in these measures, we averaged values across the six 5-min blocks following saline and CNO-2HCl injections, respectively, and compared these two averages within-subjects; testing was one-tailed because we had directional hypotheses for the changes in LFP and multi-unit measures based on our previous studies of mPFC disinhibition by picrotoxin (Pezze et al., 2014). Where the assumption of sphericity for ANOVA was violated, as indicated by a significant Mauchly’s test of sphericity, Greenhouse-Geisser correction was applied to the degrees of freedom. For pairwise comparisons of the electrophysiological average values across the 30-min periods after saline and CNO-2HCl injections, we used paired t-tests, unless the assumption of normality was violated, as indicated by a significant Shapiro-Wilk test, in which case we used nonparametric Wilcoxon signed-rank tests.

## RESULTS

### mPFC functional inhibition by muscimol, but not disinhibition by picrotoxin, impaired early reversal learning

First, we examined the effect of mPFC muscimol or picrotoxin infusion on early reversal learning (**Fig. 1A**). All infusion cannula tips were placed within the mPFC, in an area that corresponded approximately to +2.7 to +4.2 mm anterior to bregma in the atlas by Paxinos and Watson (1998) (**Fig. 1B**).

**Figure 1.**
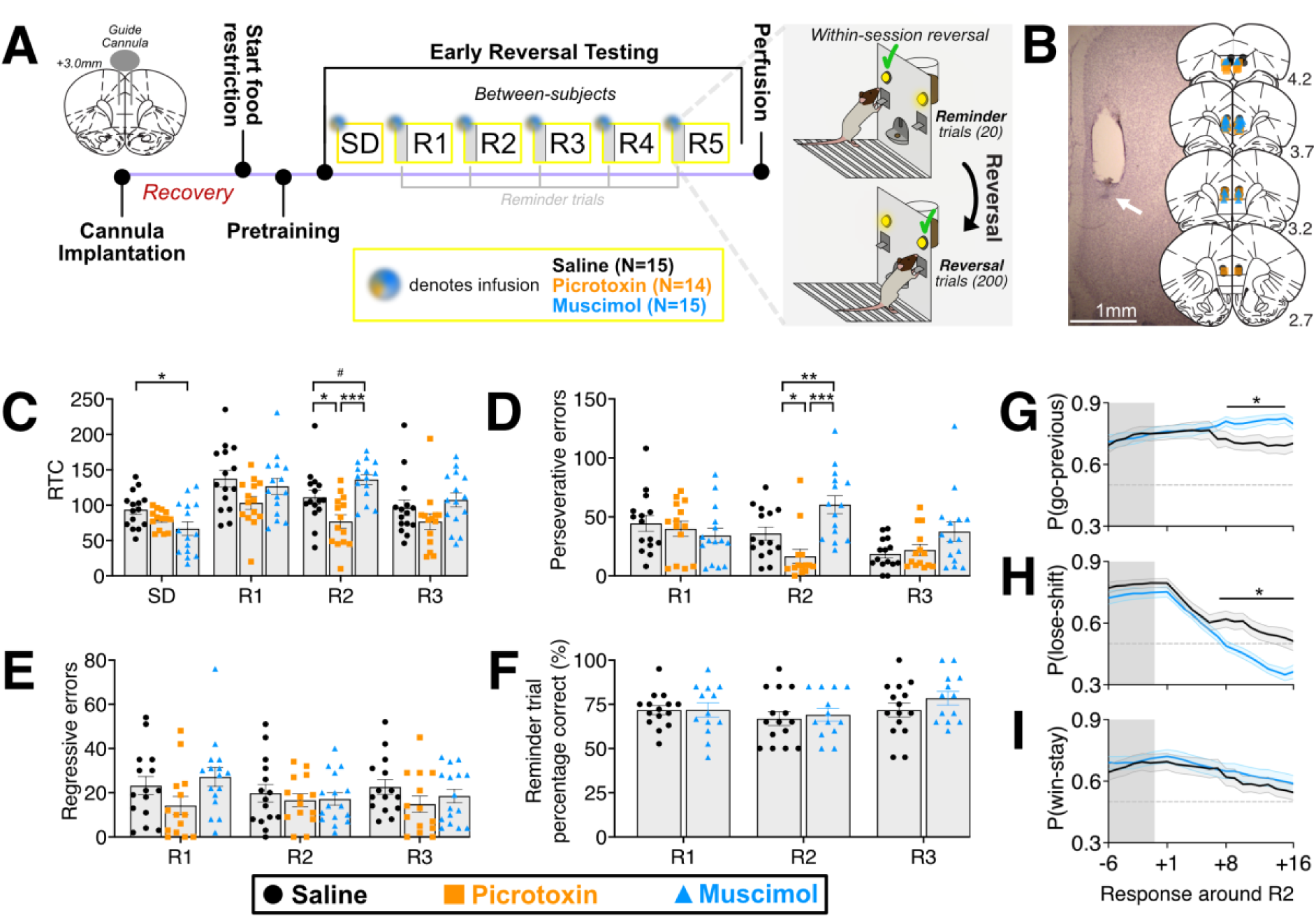
mPFC functional inhibition by muscimol, but not disinhibition by picrotoxin, impaired early reversal learning. A) Timeline of the experiment to compare the impact of mPFC infusion of saline, muscimol or picrotoxin (between-subjects) on early reversal learning, from canula implantation surgery, across food restriction, pre-training and reversal testing until transcardial perfusion. Tri-coloured dots indicate infusion time points 10 min before the spatial discrimination (SD) stage and before each reversal session at reversal stage 1 to 5 (R1 to R5). The structure of the reversal sessions is illustrated: sessions started with 20 reminder trials, during which rats were tested for their expression of the ‘old’ rule from the previous session (e.g., press the right lever), following which reward contingencies were reversed (e.g., left lever was now rewarded) and rats received a maximum of 200 reversal trials. B) Cresyl-violet-stained coronal brain section showing an infusion cannula placement in the prelimbic mPFC, with the tip indicated by white arrow. Overlayed are the approximate locations of infusion cannula tips, separated by infusion group (Saline, black dots; picrotoxin, orange squares; muscimol, blue triangles), shown on coronal plates adapted from the atlas by Paxinos and Watson (1998). Distance (in mm) anterior to bregma, according to the atlas, is indicated on the right. C-F) RTC (C), perseverative errors (D), regressive errors (E) and percentage of correct responses during reminder trials (F) in the saline, muscimol and picrotoxin groups across task stages (SD, and R1 to R3). Bar graphs show mean±SEM, with individual values for each rat plotted to show the range of data. Significant differences and trends as revealed by post-hoc pairwise comparisons are indicated: *, *p*<0.05; **, *p*<0.01; ***, *p*<0.001; #, *p*=0.051. G-I) Bayesian analysis of strategies around reversal: mPFC functional inhibition by muscimol enhanced use of old strategy (go-previous) and disrupted exploration (lose-shift) at R2. Probabilities (mean±SEM) of different strategies are shown for the 6 reminder-trial responses (grey shading) preceding reversal and the 16 responses following reversal: go-previous (G), lose-shift (H), and win-stay (I). Horizonal dashed line indicates P=0.5 (i.e., chance). Asterisks indicate responses where saline and muscimol significantly differed (p<0.05).

#### mPFC functional inhibition increased, whereas disinhibition reduced RTC during R2

Whole-session reversal performance indexed via RTC (Responses to Criterion) across the initial SD stage and the following first three reversal stages (R1 to R3) is shown in **Fig. 1C**. The saline group showed an initial reversal cost, with higher RTC at R1 compared to SD (mean±SEM, 137.47±11.97 and 93.6±6.54, respectively), which decreased across reversals as learning progressed. A significant drug × stage interaction was observed (*F*(6,123)=3.340, *p*=0.004). At R2, mPFC functional inhibition by muscimol increased RTC, whereas mPFC disinhibition by picrotoxin reduced the reversal cost (*F*(2,41)=10.892, *p*<0.001). RTC in the muscimol group were higher compared to picrotoxin (*p*<0.001) and tended to be higher compared to saline (*p*=0.051). Additionally, RTC in the picrotoxin group were lower than in the saline group (*p*=0.010). Interestingly, muscimol improved SD learning (*F*(2,41)=3.655, *p*=0.035), reducing RTC compared to saline (*p*=0.010; all other pairwise comparisons, smallest *p*=0.170). No significant differences were observed at R1 (F(2,41)=2.540, *p*=0.091) or R3 (*F*(2,41)=2.060, *p*=0.140).

#### mPFC functional inhibition increased, whereas disinhibition reduced, perseverative errors during R2

To further investigate which aspects of behavior were affected by the manipulations, we next examined perseverative and regressive errors (**Fig. 1D&E**). For perseverative errors, we observed a significant drug × stage interaction (*F*(4,82)=4.852, *p*=0.001). There was a main effect of drug at R2 (*F*(2,41)=11.625, *p*<0.001), where mPFC muscimol increased perseverative errors compared to both saline (*p*=0.009) and picrotoxin (*p*<0.001), and mPFC picrotoxin reduced perseverative errors compared to saline (*p*=0.028). In addition, a trend towards a main effect of drug was observed at R3 (*F*(2,41)=3.134, *p*=0.054), but not R1 (*F*<1). Regressive errors were similar across drug groups (*F*(2,82)=2.043, *p*=0.143) and stages (main effect of stage and drug × stage interaction, *F*<1), suggesting that mPFC manipulations specifically affected perseveration without causing non-specific learning impairments during early reversals.

#### mPFC disinhibition increased omissions, limiting reminder-trial responses

mPFC picrotoxin markedly increased the number of omissions during reminder-trial responses (*F*(2,41)=173.014, *p*<0.001), across all task stages (only a trend towards a drug × stage interaction was observed, *F*(3.432,70.365)=2.416, p=0.066), compared to saline (*p*<0.001) and muscimol (*p*<0.001), whereas saline and muscimol did not differ (*p*=0.333) (**Table S1**). Omissions across the initial SD learning trials and the reversal trials at R1-3 were also markedly increased by picrotoxin. There was a drug × stage interaction (*F*(3.895,79.856)=4.291, *p*=0.004), with a main effect of drug infusion at every stage (smallest *F*(2,41)=19.436, *p*<0.001). Specifically, at all stages, mPFC picrotoxin increased omissions compared to both saline and muscimol (both *p*<0.001), which did not differ from each other at any stage (smallest *p*=0.333). This increase in omissions by mPFC neural disinhibition is consistent with previous findings (Pezze et al., 2014; Auger et al., 2017).

#### mPFC functional inhibition did not affect correct reminder-trial responses

The marked increase in reminder trial omissions in the picrotoxin group limited the number of reminder-trial responses (average reminder-trial responses across R1-R3 out of a maximum of 20, 3.86±0.86, mean±SEM) compared to the saline (19.78±0.09) and muscimol (19.66±0.11) group. This low number of reminder trials rendered the analysis of percentage correct measures for reminder trials unreliable in the picrotoxin group. Therefore, the picrotoxin group was excluded from the analysis of the percentage correct values during reminder trials. Analysis of percentage correct measures limited to the muscimol and saline groups, revealed that prefrontal muscimol did not affect correct responses during reminder trials compared to saline (no main effect of drug or stage, and no drug × stage interaction, largest *F*(2,52)=2.149, *p*=0.127) (**Fig. 1F**).

#### mPFC disinhibition increased response latencies

Next, we examined latencies of correct and incorrect responses across reminder trials and reversal trials (**Table S1**). Due to the high number of omissions, only one rat receiving mPFC picrotoxin made at least one correct and one incorrect response during the 20 reminder trials at all three reversal stages (compared to all 15 rats in both the saline and muscimol groups). Therefore, we restricted the analysis of reminder-trial latencies across the three reversal stages to the saline and muscimol groups, which revealed no main effect or interaction involving drug (largest *F*(1,28)=1.773, smallest *p*=0.194). All three groups could be included in the analysis of latencies during the SD trials and the reversal trials of the three reversal stages (R1-3). Consistent with previous studies showing increased response latencies following mPFC disinhibition (Auger et al., 2017; Auger & Floresco, 2014; Enomoto et al., 2011; Pezze et al., 2014), this analysis revealed increased response latencies following mPFC picrotoxin regardless of response type and at all task stages. ANOVA revealed a significant drug × stage × response type (correct vs incorrect) interaction (*F*(6,117)=2.367, *p*=0.034). We further analyzed this triple interaction, using a 2-way ANOVA at each stage (SD, R1-3), with drug and response type as factors. This showed a main effect of drug at each stage (smallest *F*(2,39)=3.954, *p*=0.027), as well as a main effect of response type at SD, R1 and R3 (smallest *F*(1,39)=7.652, *p*=0.009). Furthermore, there was a significant drug × response type interaction at R1 (*F*(2,39)=6.364, *p*=0.004), but not the other stages (largest *F*(2,39)=1.191, *p*=0.315). *Post-hoc* pairwise comparisons at SD, R2 and R3 revealed that, at all these stages, picrotoxin increased response latencies compared to saline and muscimol infusions (largest *p*=0.008), which did not differ (smallest *p*=0.157). Simple main effects analysis of the interaction at R1 revealed a main effect of drug on both correct and incorrect latencies (both *F*(2,39)>12.8, *p*<0.001). *Post-hoc* pairwise comparisons revealed that, also at R1, picrotoxin increased both correct and incorrect response latencies compared to saline and muscimol (largest *p*=0.004), which did not differ from each other (smallest *p*=0.333).

#### Bayesian analysis of strategies around reversal: mPFC functional inhibition enhanced use of the old strategy (go-previous) and disrupted exploration (lose-shift) at R2

To complement classical whole-session measures of reversal performance, and to gain an insight into the underlying behavioral changes across the trials within a session, we first examined rats’ response strategies during responses around rule reversal (6 reminder-trial responses and 16 reversal-trial responses), using the Bayesian trial-by-trial strategy analysis (Maggi et al., 2024). The picrotoxin group was excluded from this analysis, because of the low number of reminder trials completed by this group due to the high number of omissions (see above). Analysis of the go-previous probability in the saline and muscimol groups across all reversal stages (R1-3, data not shown) revealed a significant drug × stage × response interaction (*F*(42,1176)=2.235, *p*<0.001). Further analysis of this triple interaction revealed a significant drug × response interaction at R2 (*F*(3.340,93.078)=6.456, *p*<0.001) (**Fig. 1G**), whereas, at R1 and R3, there was no main effect or interaction involving drug (largest *F*(2.461,68.910)=1.350, *p=*0.267) (data not shown). The drug × response interaction at R2 reflected that mPFC functional inhibition increased the go-previous probability compared to saline between responses 9-15 after reversal (smallest *F*(1,28)=5.165, *p*=0.031), with trends toward an increase at responses 8 (*F*(1,28)=3.980, *p*=0.056) and 16 (*F*(1,28)=4.090, *p*=0.053) after reversal, in line with the increased RTC and perseverative errors caused by mPFC muscimol at that stage (**Fig. 1C&D**).

Before rule reversal, during reminder trials, rats showed high lose-shift (**Fig. 1H**) and high win-stay (**Fig. 1I**) probabilities. The high lose-shift probabilities may partly reflect that rats made very few incorrect responses during reminder trials, and, therefore, the lose-shift probability was hardly updated by our Bayesian analysis algorithm and remained at similarly high values as at the end of the preceding reversal session. The high win-stay probabilities indicated strong exploitation, consistent with the high percentage of correct responses during reminder trials. However, immediately after reversal, there was a steep decline in lose-shift behavior and a more gradual decline in win-stay behavior, similar to our previous findings following the switch from a spatial rule to a cue-based rule (Maggi et al., 2024), revealing that the rule reversal induced marked changes in exploitation and exploration. Analysis of lose-shift probabilities across R1 to 3 (data not shown) revealed a drug × stage × response interaction (*F*(42,1176)=1.436, *p*=0.037). This triple interaction reflected that lose-shift behavior was reduced by muscimol, compared to saline, specifically at R2, between responses 8 to 16 after reversal (drug × response interaction, *F*(1.803,50.471)=3.569, *p*=0.040; main effect of drug at responses 8-16, smallest *F*(1,28)=6.795, *p*=0.014; main effect of drug for all other responses, largest *F*(1,28)=2.797*, p*=0.106) (**Fig. 1H**). In contrast, at R1 and R3, there was no main effect or interaction involving drug (largest *F*(1,28)=1.250, *p*=0.273), only a main effect of response (both *F*(2.094,54.390)>40, *p*<0.001) (data not shown). In contrast to lose-shift behaviour, win-stay behaviour was not affected by mPFC drug infusions, but declined similarly following both mPFC saline and muscimol and across all reversal stages following reversal (only a main effect of response, *F*(2.863,80.174)=81.264, *p*<0.001, but no other main effects or interactions, largest *F*(42,1176)=1.289, *p*=0.104) (data for R2 are shown in **Fig. 1I**).

*Bayesian analysis of strategies preceding learning: no effect of mPFC functional inhibition* Contrasting with the group differences in strategy use around reversal, mPFC muscimol did not affect strategies leading up to the point of learning (data not shown). Analysis of go-new, lose-shift and win-stay probabilities during responses −5 to −1 leading up to learning only revealed an effect of response for each strategy (smallest *F*(1.223,34.246)=5.288, *p*=0.022), reflecting that these task-pertinent strategies increased leading up to reaching the performance criterion, without any other main effect or interaction involving drug (largest *F*(1.387,38.830)=1.667, *p*=0.203).

### mPFC disinhibition by picrotoxin, but not functional inhibition by muscimol, impaired serial reversal learning

Next, we tested the effect of mPFC muscimol or picrotoxin infusion on serial reversal learning, when reversal performance was well-established, across R5 to 10 (**Fig. 2A**). All infusion cannula tips were placed within the mPFC (**Fig. 2B**).

**Figure 2.**
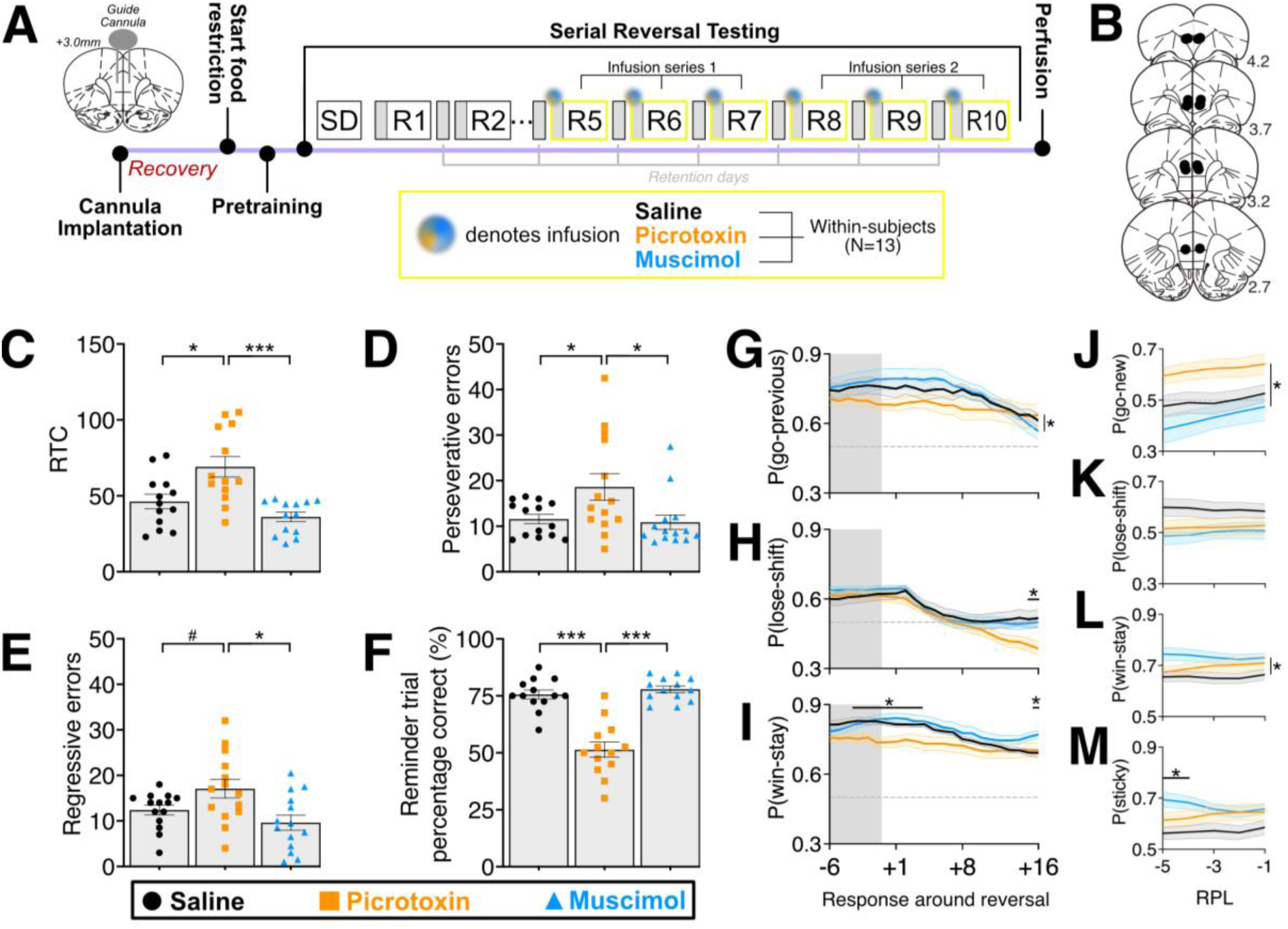
mPFC disinhibition by picrotoxin, but not functional inhibition by muscimol, impaired serial reversal learning. A) Timeline of the experiment to examine the impact of mPFC infusion of saline, muscimol or picrotoxin (within-subjects) on serial reversal learning, from cannula implantation surgery, across food restriction (F.R), spatial discrimination (SD) and early reversal (reversals 1 to 4, R1 to 4) training and serial reversal testing (R5 to R10) until transcardial perfusion. Tri-coloured dots indicate infusion time points 10 min before R1 to R5. Retention days were included between reversal sessions, and rats completed two infusion series, where they underwent the same series of three different infusions (saline, muscimol and picrotoxin) twice. B) Approximate locations of infusion cannula tips (black dots), shown on coronal plates adapted from the atlas by Paxinos and Watson (1998). Distance (in mm) anterior to bregma, according to the atlas, is indicated on the right. C-F) RTC (C), perseverative errors (D), regressive errors (E) and percentage of correct responses during reminder trials (F) in the saline, muscimol and picrotoxin conditions, averaged across both infusion series. Bar graphs show mean±SEM, with individual values for each rat plotted to show the range of data. Significant differences as revealed by post-hoc pairwise comparisons are indicated: *, *p*<0.05; **, *p*<0.01; ***, *p*<0.001. G-I) Probabilities (mean±SEM) of different strategies are shown for the 6 reminder-trial responses (grey shading) preceding reversal and the 16 responses following reversal: go-previous (G), lose-shift (H), and win-stay (I). Horizonal dashed line indicates P=0.5 (i.e., chance). Asterisks above datapoints indicate responses where groups significantly differed (p<0.05) following observation of a drug × response interaction. Main effect of drug across responses, without an interaction, is indicated by a horizontal line and asterisk to the right of the data lines. J-M) Probabilities (mean±SEM) of different strategies are shown for the 5 responses preceding learning (RPL): go-new (J), lose-shift (K), win-stay (L), and sticky (M). Horizonal dashed line indicates P=0.5 (i.e., chance). Significant differences are indicated as for G-I.

#### mPFC disinhibition increased RTC during serial reversal learning

mPFC picrotoxin markedly impaired serial reversal performance, reflected by higher RTC (main effect of drug, *F*(2,24)=9.603, p<0.001), compared to saline (*p*=0.003) and muscimol (*p=*0.003), which did not differ from each other (*p*=0.266) (**Fig 2C**). There was a trend towards a main effect of infusion series (*F*(1,12)=4.255, *p*=0.058), but no drug × infusion series interaction (*F*<1), which reflected that RTC tended to decline slightly from series 1 to 2 across all infusion conditions (data not shown).

#### mPFC disinhibition increased both perseverative and regressive errors

The increase in RTC by mPFC picrotoxin reflected an increase in perseverative errors (main effect of drug, *F*(1.269,15.231)=4.592, *p*=0.041) compared to saline (p=0.021) and muscimol (*p*=0.011) **(Fig. 2D)**, and in regressive errors (main effect of drug, *F*(2,24)=5.427, *p*=0.011) compared to muscimol (*p*=0.003), with a trend toward an increase compared to saline (*p*=0.050) (**Fig. 2E**). Errors did not differ between saline and muscimol conditions (smallest *p*=0.333). There was no main effect of infusion series or drug × infusion series interaction (largest *F*(1.199,14.383)=1.199, *p*=0.140).

#### mPFC disinhibition impaired expression of the previous rule during reminder trials

mPFC disinhibition markedly impaired the expression of the old rule prior to rule change, as reflected by a reduced percentage of correct responses during reminder trials ([correct responses/all responses]X100%) (main effect of drug, F(2,24)=24.673, *p*<0.001; **Fig 2F**). The percentage of correct reminder trials was reduced by picrotoxin compared to saline and muscimol infusions (both *p*<0.001), which did not differ from each other *(p=*0.333). No main effect or interaction involving infusion series was observed (main effect of infusion series, *F*<1; drug × infusion series interaction, *F*(1.191,22.939)=2.778, *p*=0.085).

#### mPFC disinhibition increased omissions and response latencies

Consistent with previous studies and our early reversal learning experiment (see above), mPFC disinhibition markedly increased omissions and latencies of correct and incorrect responses across both reminder and reversal trials, whereas functional inhibition did not affect these measures (**Table S2**). However, compared to our early reversal learning experiment, omissions were overall lower, and the increase in omissions by mPFC picrotoxin was not as detrimental to reminder trial responding. Nevertheless, a main effect of drug was still observed for reminder-trial omissions (*F*(1.003,12.04)=5.510, *p*=0.037), and reversal-trial omissions (*F*(1.002,12.02)=24.204, *p*<0.001); in both cases, this was driven by increased omissions following mPFC picrotoxin compared to saline and muscimol (largest *p*=0.008), whereas saline and muscimol did not differ (both *p*=0.333). No main effect or interaction involving series was observed for omissions during reminder or reversal trials (largest *F*(1,12)=2.695, *p*=0.127). For reminder-trial response latencies, there was a significant drug × response-type interaction (*F*(2,22)=4.317, *p*=0.026), and no other main effect or interaction involving infusion series or response type (all *F*<1). Simple-main effects analysis showed a significant effect of drug for both correct- and incorrect-response latencies (smallest *F*(2,24)=7.279, *p*=0.004). Pairwise comparisons revealed that picrotoxin increased both correct and incorrect reminder-trial response latencies compared to saline and muscimol (all *p*<0.001), which did not differ (*p*=0.333). For reversal-trial response latencies, there was also a significant drug × response-type interaction (*F*(2,24)=3.902, *p*=0.034), alongside a significant drug × infusion series interaction (*F*(1.078,12.940)=4.747, *p*=0.046). Subsequent simple main effects analyses revealed significant main effects of drug across infusion series and response types (smallest *F*(2,24)=28.624, *p*<0.001). Pairwise comparisons revealed that this effect was driven by increased response latencies following picrotoxin, compared to saline and muscimol for both series and response types (largest *p*<0.001); muscimol and saline did not differ regardless of infusion series or response type (all *p*=0.333).

#### Bayesian analysis of strategies around reversal: mPFC disinhibition reduced adherence to old strategy (go-previous), as well as exploitation (win-stay) and exploration (lose-shift)

During serial reversals, rats receiving mPFC saline or muscimol infusions showed high go-previous probabilities during the 6 last responses before the rule change; go-previous probabilities remained high until about the 7th response after rule change when they began to decrease (**Fig. 2G**), indicating that rats abandoned the previous rule and switched to the opposite lever. This decline in go-previous behavior was evident regardless of infusion series (i.e., during R5 to 7 and R8 to 10) and differed from the pattern we observed during early reversals (R1 to R3), where the go-previous probability did not decline across the first 16 responses following rule reversal (**Fig. 1G**). Following mPFC disinhibition, however, the pattern differed: the go-previous probability tended to be reduced compared to saline and muscimol infusions during the reminder trials, at or below P=0.7 (in line with reduced reminder trial accuracy, **Fig. 2F**), but hardly declined following rule reversal, such that, towards the end of the first 16 responses after reversal, the go-previous probability was similar in all three infusion conditions (**Fig. 2G**). In support of these findings, ANOVA of the go-previous probabilities revealed a drug × response interaction (*F*(42,504)=1.651, *p*=0.008) and a trend towards a drug × infusion series interaction (*F*(2,24)=2.965, *p*=0.071), but no drug × infusion series × response interaction (*F*(42,504)=1.213, *p*=0.174). Simple main effects analysis of the drug × response interaction did not indicate a significant effect of drug for any response but indicated trends toward such an effect for responses −1, +2, and +8 around reversal (smallest *F*(2,24)=2.586, *p*=0.096); for all other responses, largest *F*(2,24)=2.542, *p*=0.100).

The time course of lose-shift behavior during repeated reversals (**Fig. 2H**) was similar to that during early reversals stages (**Fig. 1H**). Before reversal, lose-shift probability was high across all three infusion conditions, and, after reversal, there was a steep decline in this strategy, indicating a tendency to stick with the previous, but now incorrect, response for several trials. Following mPFC saline and muscimol infusions, this decline plateaued around 5 responses after reversal, coinciding with a decline in go-previous probability (**Fig. 2G**), which reflected that rats began to switch to the opposite lever. However, following mPFC picrotoxin, lose-shift probability declined throughout all 16 responses that were analysed following reversal, dropping below the lose-shift probabilities in the saline and muscimol conditions at response 15 and 16. Importantly, the similar lose-shift probabilities in all three infusion conditions prior to response 15 post-reversal suggest that mPFC picrotoxin does not reduce lose-shift probabilites beyond the typical decline seen following an unexpected rule change. Rather, mPFC picrotoxin appears to impair the mechanisms underlying prompt strategy adjustment, resulting in failure to rescue appropriate lose-shift behavior over time. These findings were supported by ANOVA of lose-shift probabilities, which revealed a significant drug × response interaction (*F*(42,504)=2.250, *p*<0.001), with no further main effects or interactions (largest *F*(42,504)=1.096, *p*=0.319). Simple main effect analysis revealed a significant effect of drug at responses 15 and 16 (smallest *F*(2,24)=4.185, *p*=0.028) and a trend at responses 13 and 14 post-reversal (smallest *F*(2,24)=3.185, *p*=0.059) (all other responses: largest *F*(2,24)=2.285, *p*=0.123). *Post-hoc* pairwise comparisons showed that picrotoxin reduced lose-shift probabilities at both responses 15 and 16 compared to saline (largest *p*=0.043) and muscimol (largest *p*=0.029), which did not differ significantly from each other (*p*=0.600).

The probability of win-stay behaviour during serial reversals remained high around reversal, contrasting with the steep decline following reversal during early reversals stages (**Fig. 1I**), but was reduced by mPFC picrotoxin compared to saline and muscimol during some of the responses around reversal (**Fig. 2I**). ANOVA of win-stay probabilities revealed a significant drug × response interaction (*F*(42,504)=2.904, *p*<0.001), with no further main effects or interactions (largest *F*(21,252)=1.196, *p*=0.254). Simple main effect analyses revealed a significant drug effect starting four responses before, and ending four responses after reversal, as well as at the 16th response after reversal (smallest *F*(2,24)=3.778, *p=*0.037). Additionally, there was a trend toward an effect of drug at the fifth response before and after reversal, as well as at the 15th response after reversal (smallest *F*(2,24)=2.591, *p=*0.096) but not drug effect at any other response (largest *F*(2,24)=2.311 *p*=0.121). *Post-hoc* pairwise comparisons, to analyze further the simple main effects of drug at responses −4 to +4 around reversal, showed that prefrontal picrotoxin significantly reduced win-stay probability compared to saline and muscimol (largest *p*=0.049), which did not differ (smallest *p*=0.260). On the other hand, the simple main effect of drug at response 16 after reversal was driven by increased win-stay probability in the muscimol condition, compared to saline (*p*=0.019) and picrotoxin (*p*=0.036), which did not differ between each other at that stage (*p*=0.333).

#### Bayesian analysis of strategies preceding learning: mPFC disinhibition increased adherence to new rule (go-new) and functional inhibition increased win-stay and sticky behavior leading up to learning

Across the 5 responses leading up to the point of learning, the go-new probabilities were increased by mPFC picrotoxin compared to saline and muscimol infusions and gradually increased regardless of the infusion condition (**Fig. 2J**). The increased go-new probability, alongside increased RTC (**Fig. 2C**), suggests that, following mPFC picrotoxin, rats achieved the learning ‘breakthrough’ (i.e., 10 consecutive correct responses) at a later stage (as reflected by higher RTC) because they required prolonged accumulation of evidence for ‘go-new’ (i.e., P(go-new)>0.5) before reaching the criterion. ANOVA of go-new probabilities revealed a main effect of drug (*F*(2,24)=4.650, *p*=0.020) and of response (*F(*1.526, 18.318)=16.264, *p*<0.001), reflecting a gradual increase, alongside a trend towards an effect of series (*F*(1, 12)=3.718, *p*=0.078), but no interactions (largest *F*(2.112,25.343)=1.165, *p*=0.330). *Post-hoc* pairwise comparisons showed that, following picrotoxin infusions, go-new probabilities were higher than following muscimol infusions (*p*=0.006) and tended to be higher compared to saline infusions (*p*=0.063), with no differences between the latter (*p*=0.304).

Lose-shift probabilities were relatively stable during the last 5 responses preceding learning and were numerically higher in the saline condition (**Fig. 2K**), but this difference was not significant, with ANOVA not revealing any main effect or interaction involving drug (largest *F*(3.139, 37.664)=2.180, *p*=0.104).

However, mPFC muscimol increased the probability of win-stay behavior (**Fig. 2L**) and of sticky behavior compared to saline and picrotoxin (**Fig. 2M**). ANOVA of win-stay probabilities revealed a main effect of drug (*F*(2,24)=5.922, *p*=0.008), alongside a trend for an effect of infusion series (*F*(1,12)=4.265, *p*=0.061) (all other main effects or interactions: largest *F*(2.534,30.408)=1.874, *p*=0.162). *Post-hoc* pairwise comparisons showed that the win-stay probabilities were higher following muscimol compared to saline (*p*=0.002) and tended to be higher compared to picrotoxin (*p*=0.095). Saline and picrotoxin did not differ (*p*=0.101). ANOVA of sticky behavior revealed a drug × response interaction (*F*(2.849,34.186)=3.354, *p*=0.032), but no other main effects or interactions (largest *F*(1,12)=2.757, *p*=0.123). Simple main effects analysis revealed an effect of drug at responses −5 and −4 before reaching criterion (smallest *F*(2,24)=4.555, *p*=0.021) and trends at responses −3 and −2 (largest *F*(2,24)=2.902, *p*=0.074), but no drug effect at response −1 (*F*(2.24)=2.012, *p*=0.156). *Post-hoc* pairwise comparisons indicated higher sticky probability in the muscimol condition compared to saline and picrotoxin at response −5 (largest *p*=0.044) and to saline (*p*=0.006), but not picrotoxin (*p*=0.119), at response −4. Saline and picrotoxin did not differ from one another at any point (smallest *p*=0.175). This suggests that, following mPFC muscimol, rats reached criterion during serial reversal learning by using slightly different strategies than in the other two conditions.

### Chemogenetic mPFC disinhibition: Characterisation of hM4Di DREADD expression and functionality in mPFC GABA neurons of Vgat-Cre rats

#### Histological verification of spread, penetrance and specificity of hM4Di expression in Vgat-Cre rats

We used an initial cohort of Vgat-Cre rats (n=6; **Table 1**, Cohort #3) rats to characterize DREADD expression in the mPFC. Visualisation of the endogenous mCherry tag of DREADD receptors in the brains of these rats revealed bilateral DREADD expression throughout the mPFC (**Fig. 3A**). In most rats, expression was restricted to an area corresponding to approximately +4.7 to +2.2 mm anterior to bregma according to the atlas by Paxinos and Watson (1998). There was some DREADD expression outside the mPFC, primarily within the motor cortex (primary and secondary) between +4.2 and +2.7 mm from bregma, and in two cases extending to early ventral and dorsal orbitofrontal areas at +5.2 mm from bregma.

**Figure 3.**
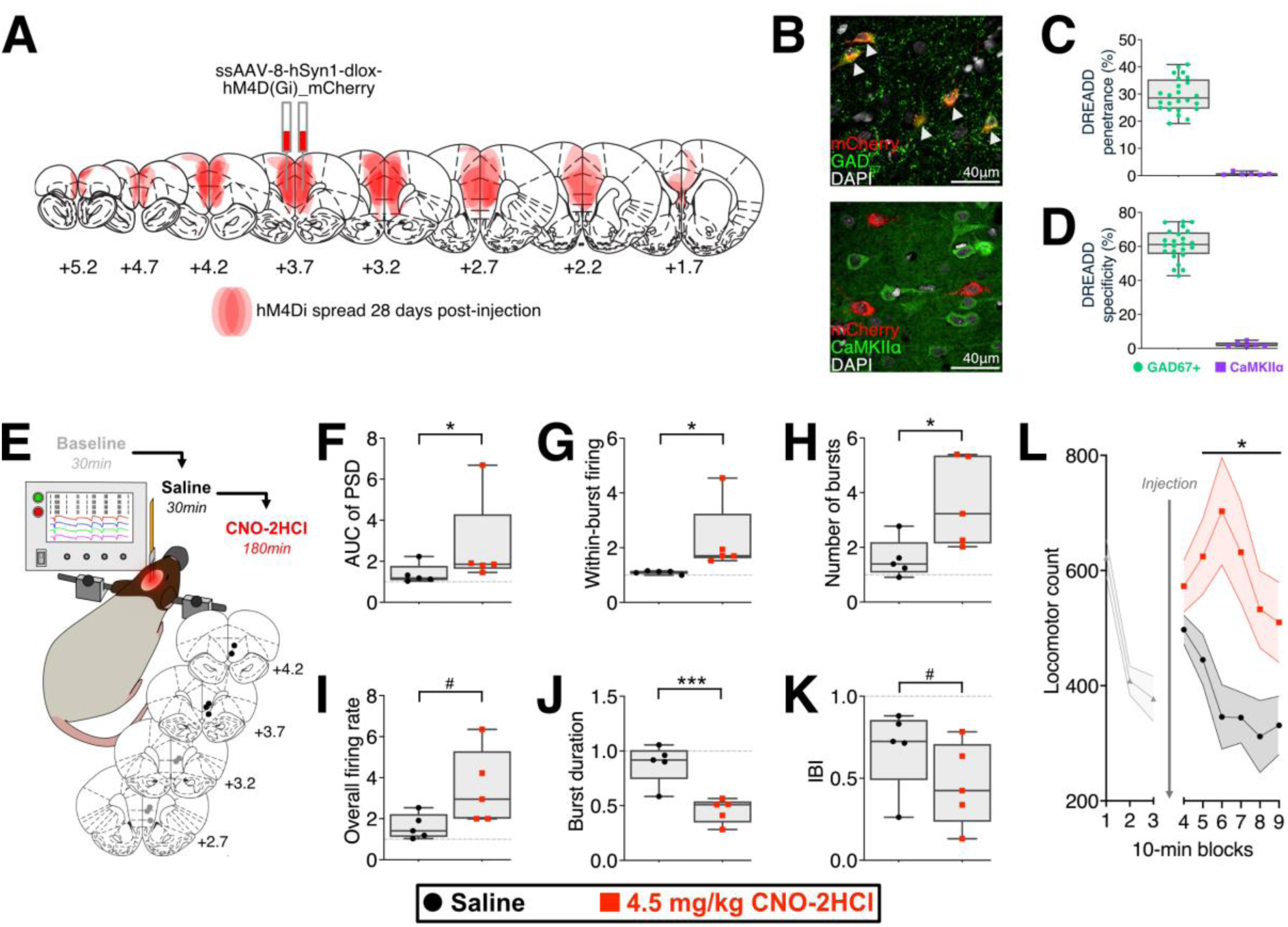
Chemogenetic mPFC disinhibition: Characterisation of mPFC hM4Di DREADD expression and functionality in Vgat-Cre rats. A) DREADD (mCherry) spread across mPFC following viral vector injection at +3.2 mm. For each individual rat, the area showing mCherry signal was uniformly shaded (red) on coronal sections taken from the brain atlas by Paxinos and Watson (1998). The shadings were overlaid for all rats (n=6), meaning darker red areas indicate overlap of DREADD expression across rats. Distance from bregma (in mm), according to the atlas, is indicated underneath each section. B) Image showing staining for GAD67 (top; green), a marker of GABAergic neurons, and CaMKIIa (bottom; green), a marker of excitatory neurons, alongside mCherry signal (red) indicating hM4Di expression. White arrow tips highlight mCherry (hM4Di) expression in GABAergic cells. In both images, nuclear DAPI staining (white) was used to indicate overall cell count. C) Box and whisker plot (Whiskers: min-max; Hinges: 25^th^-75 percentile; Line: median) of DREADD penetrance in GAD67+ and CaMKIIa+ cells (i.e., proportion of cells of this type that expressed mCherry, the DREADD tag) in slices taken close to injection site (around +3.2 mm from bregma). Individual dots represent values obtained from a single slice. We used four slices per rat for GAD67 (resulting in 24 values for 6 rats) (green circles) and one slice for CaMKIIa (resulting in 6 values for 6 rats) (red squares), as the latter was predominantly a control staining to ensure no expression in excitatory cells. D) Box and whisker plot of cell-type specificity of DREADD expression (proportion of total mCherry signal attributable to each neuronal subtype). For explanation of the values shown, see legend to C). E) The top panel illustrates the in vivo electrophysiological procedure. Rats were anesthetised, fixed in a stereotaxic frame, and electrode arrays consisting of four microwire implanted unilaterally into the mPFC. Recording began with a 30-min baseline period. This was followed by an i.p. saline injection and another 30-min recording period. After this, CNO-2HCl (4.5 mg/kg; i.p.) was injected followed by a 180-min recording period. Following recording, the most anterior and most posterior electrode tips were electrolytically marked, and rats were transcardially perfused, and brains extracted. The overlapping bottom panel shows approximate locations of electrode tips (most anterior: grey; most posterior: black) on coronal plates adapted from the atlas by Paxinos and Watson (1998). Distance (in mm) anterior to bregma is indicated on the left, according to the atlas. F-K) Chemogenetic mPFC disinhibition enhanced mPFC neural burst firing. Box and whisker plots (Whiskers: min-max; Hinges: 25^th^-75 percentile; Line: median) show measures of LFP (AUC of PSD) and multi-unit activity (within-burst firing rate, number of bursts, overall firing rate, burst duration and inter-burst interval -IBI-, respectively) following systemic saline (black circles) and CNO-2HCl (4.5 mg/kg; red squares), within-subjects. All values are normalized to baseline. Datapoints for each rat were averaged across the first 6 5-min blocks following saline and CNO-2HCl injection, respectively (30 min post-injection). Significant differences and trends revealed by Wilcoxon signed rank test (one-tailed) are indicated: *, *p*<0.05; ***, *p*<0.001; #, 0.05 <*p*>0.1. L) Chemogenetic mPFC disinhibition increased locomotor activity. Locomotor counts (mean±SEM) are shown for an initial 30-min baseline period (grey triangles) prior to injection (values are averaged across days, as no significant difference in baseline activity was observed *F*<1) and for the 60-min period following injection of saline (black circles) or CNO-2HCl(4.5 mg/kg; red squares)(n=12 rats, within-subjects design). Asterisk and line indicate the 10-min blocks where saline and CNO-2HCl conditions significantly differed (p<0.05).

Immunohistochemical staining of the brains of the same rats revealed good DREADD penetrance in GABA neurons (GAD67+ cells) and specificity of expression to GAD67+ cells, compared to cells labelled with the excitatory cell marker CaMKIIa+ (**Fig. 3B, C, D**). As to penetrance, 29.58±1.23% (mean±SEM) of all GAD_67_+ cells within the slice most proximal to the injection site (i.e., around +3.2 mm from bregma) expressed the DREADD, compared to 0.67±0.19% of all CaMKIIα cells (**Fig. 3C**). As to cell-type specificity, 60.8±1.81% (**Fig. 3D**) of all DREADD expression was localised to GAD_67_+ cells, whereas only 2.35±0.52% was localised in CaMKIIa+ cells. Inspection of CaMKIIa co-localisation revealed that proximity of mCherry-expressing GABA+ cells to CaMKIIa+ cells led to the software misclassifying several CaMKIIa+ cells as mCherry+, driving the negligible mCherry-CaMKIIa colocalization. In contrast, GABA and mCherry classification was not affected by this, as overlap for these markers was substantial, reflecting true co-localisation. Finally, there was some co-expression of mCherry in ChAT+ cells (not shown), with an average co-localisation of 5.33±0.89%, which, upon closer inspection, was true mCherry expression in ChAT+ cells, probably reflecting a sub-population of GABA-acetylcholine co-releasing neurons (Granger et al., 2016; Saunders et al., 2015). The remaining difference between total DREADD expression, i.e. the difference between 100% and all cell-type specific expression (adding up to only about 70%), likely reflects underestimation of GABA cells, due to either incomplete staining or false-negative classification by the analysis software. Overall, our histological analyses revealed good penetrance and cell-type specificity, consistent with values reported in previous studies using cell-type specific DREADD expression (Nguyen et al., 2014; Smith et al., 2016).

#### Chemogenetic mPFC disinhibition enhanced mPFC neural burst firing

The neural effects of activating the DREADD receptors expressed within the mPFC were initially examined using 6.0 mg/kg CNO-2HCl (n=5, **Table 1**, Cohort #3). A notable intensification in burst firing and increased LFP power were observed (**Fig. S1**). However, our reversal study indicated that 6.0 mg/kg caused non-specific behavioral effects, highlighting the unsuitability of that dose, and precluding further analysis (data not shown). Therefore, in a subset of the rats that had completed the reversal (and locomotor testing, see below) (n=5; **Table 1**, Cohort #4) we re-examined the effects of CNO-2HCl on mPFC neural activity at a dose of 4.5 mg/kg, which produced reversal and locomotor changes without gross non-specific effects. All electrodes were included in the analysis and were placed within the mPFC between +4.7 and +2.7 mm anterior to bregma (**Fig 3E**), surrounded by strong DREADD expression (see **Fig 4B**).

**Figure 4.**
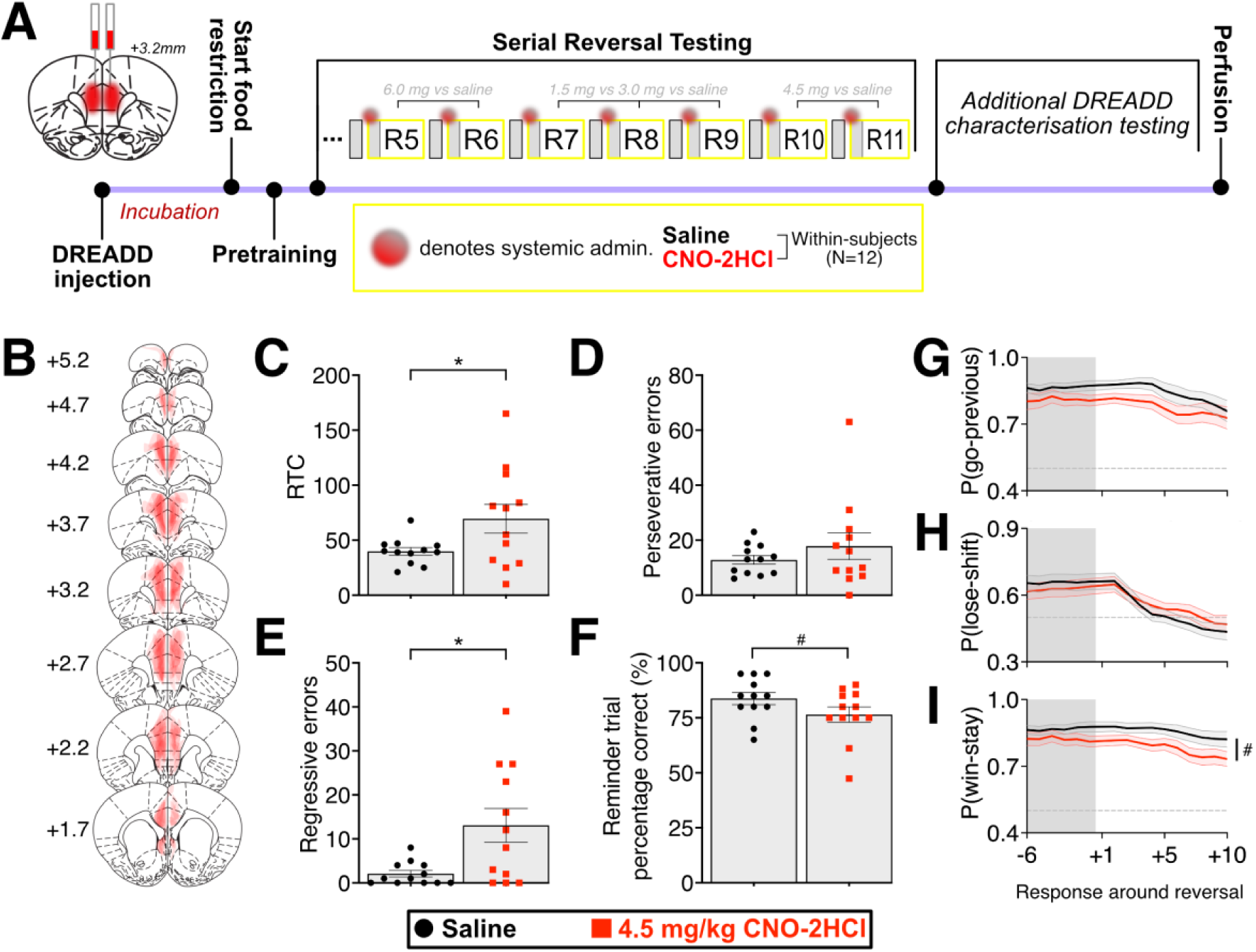
Chemogenetic mPFC disinhibition impaired serial reversal learning. A) Timeline of the experiment examining the impact of chemogenetic mPFC disinhibition on serial reversal learning. Vgat-Cre rats (n=12) underwent surgery to inject the inhibitory hM4Di DREADD into the mPFC, followed by 28-day incubation period to allow for expression of the DREADD in GABAergic mPFC neurons, before undergoing food restriction and pre-training on the operant reversal task up to reversal 4. The impact of CNO-2HCl and saline injection was then compared within-subjects across reversals 5 to 11 (R5-11). Several doses of CNO-2HCl were tested, but only the findings following 4.5 mg/kg are shown in this figure. Following reversal testing, rats were used for additional characterisation of DREADD functionality in the mPFC (locomotor activity n=12; in vivo electrophysiology n=5; see Fig. 3). B) DREADD (mCherry) spread across mPFC. For each individual rat, the area showing mCherry signal was uniformly shaded (red) on coronal sections taken from the brain atlas by Paxinos and Watson (1998). The shadings were overlaid for all rats, meaning darker red areas indicate overlap of DREADD expression across rats. Distance from bregma (in mm), according to the atlas, is indicated on the left. C-F) RTC (C), perseverative errors (D), regressive errors (E) and percentage of correct responses during reminder trials (F) following systemic injection of saline (black circles), or CNO-2HCl (4.5 m/kg; red squares). Bar graphs show mean±SEM, with individual values for each rat plotted to show the range of data. Significant differences and trends as revealed by post-hoc pairwise comparisons are indicated: *, *p*<0.05; #, p=0.078. G-I) Probabilities (mean±SEM) of different strategies are shown for the 6 reminder-trial responses (grey shading) preceding reversal and the 10 responses following reversal: go-previous (G), lose-shift (H), and win-stay (I). Horizonal dashed line indicates P=0.5 (i.e., chance). The trend towards a main effect of drug for win-stay is indicated by a horizontal line and ‘#’ next to data lines (p=0.087).

Similar to our previous findings following mPFC picrotoxin (Pezze et al., 2014), chemogenetic disinhibition of the mPFC by 4.5 mg/kg of CNO-2HCl increased LFP power, measured as AUC of PSD (W=0, z=-2.023, p=0.031, **Fig. 3F)**, and intensified neuronal burst firing, reflected by increased within-burst firing rate (W=0, z=-2.023, p=0.031, **Fig. 3G)** and number of bursts per 5-min block (t(4)=2.807, p=0.024, **Fig. 3I**); overall firing rate also tended to be increased (W=1, z=-1.753, p=0.063, **Fig. 3H**). Additionally, a reduction in burst duration (t(4)=9.848, p<0.001, **Fig. 3J)** and trend for a reduced inter-burst interval (t(4)=2.017, p=0.057, **Fig. 3K**) was also observed. Overall, our in vivo electrophysiological measurements support that chemogenetic mPFC disinhibition causes neural effects, especially enhanced burst firing, similar to mPFC disinhibition by picrotoxin (Pezze et al., 2014).

To assess reproducibility of mPFC LFP and multi-unit measures across our studies, we compared baseline values (mean±SEM) of these measures recorded from isofluorane-anaesthetised Vgat-Cre rats in the present study to those that we recorded previously in isofluorane-anaesthetised Lister hooded rats, as part of our studies of the effects of mPFC drug microinfusions (Pezze et al., 2014). In the previous study, we had baseline values from three different prospective mPFC microinfusion groups, and we averaged these baseline values across these three groups for the purpose of the present comparison. Within-burst firing rates, number of bursts within a 5-min block, mean burst duration, and inter-burst interval were comparable between current (181.57±26.23 spikes/s, 134.3±5.86, 223.0±47.0 ms, and 5.37±1.46 s respectively) and previous work (187.15±24.35 spikes/s, 132.2±35.76, 307.5±53.0 ms, and 6.625±0.44 s respectively). In contrast, baseline overall firing rate and AUC of PSD in the present study were lower compared to our previous work (11.54±2.21 vs. 20.68±5.65 spikes/s and 0.007±0.0007 µV^2^ vs. 0.015±0.004 µV^2^, respectively). Overall, these comparisons indicate good agreement of mPFC recordings across our two studies. As to the numerical differences noted for some measures, it is unclear whether these reflect strain differences or methodological differences (including an indwelling infusion cannula in the previous study and different microwire arrays).

#### Chemogenetic mPFC disinhibition increased locomotor activity

In the Vgat-Cre rats used in the main reversal study (n=12; **Table 1**, Cohort #4), we also examined the locomotor effects of chemogenetic mPFC disinhibition by 4.5 mg/kg CNO-2HCl. Consistent with the locomotor hyperactivity caused by mPFC disinhibition by picrotoxin (Pezze et al., 2014), chemogenetic disinhibition of the mPFC increased open-field locomotor activity (**Fig 3L**). ANOVA of post-injection locomotor counts revealed a significant drug × 10-min block interaction (F(2.837, 31.203)=3.659, p=0.025). Simple main effects analysis of the drug effect at the different 10-min blocks following injection revealed that, compared to saline, CNO-2HCl injection increased locomotor activity across most of the 60 min post-injection (lowest *F*(1,11)=6.901, *p*=0.024) period, except for the first 10-min point (*F*(1,11)=2.711, *p*=0.128). As expected, locomotor counts during the three 10-min baseline blocks prior to injection did not differ between injection conditions (main effect or interaction involving injection, *F*<1); there was only a main effect of 10-min block (*F*(1.353,14.883)=38.435, *p*<0.001), reflecting locomotor habituation. Finally, there was no carry over effect of mPFC chemogenetic disinhibition on locomotor activity during the 30-min test sessions on the days following the injection days (main effect or interaction involving drug, both *F*<1), with only the main effect of time persisting (*F*(2.174,23.917)=56.397, *p*<0.001; data not shown).

### Chemogenetic mPFC disinhibition impaired serial reversal learning

#### Chemogenetic mPFC disinhibition increased RTC and regressive errors, but not perseverative errors

Complementing our studies examining the impact of mPFC muscimol and picrotoxin on reversal learning, we aimed to test if chemogenetic disinhibition of the mPFC would reproduce the marked serial reversal learning impairment caused by mPFC disinhibition by picrotoxin (**Fig 4A**). In all rats (n=12; **Table 1**, Cohort #4), the DREADD was expressed within the mPFC, without substantial expression outside the mPFC (**Fig 4B**). Chemogenetic disinhibition of the mPFC during serial reversal learning, by injection of 4.5 mg/kg CNO-2HCl, increased RTCs compared to saline injection (*F*(1,11)=6.638, *p*=0.026; **Fig. 4C**). Interestingly, chemogenetic mPFC disinhibition did not affect perseverative errors (*F*<1; **Fig. 4D**), contrasting with the increase of perseverative errors by mPFC picrotoxin (**Fig. 2D**), but only increased regressive errors (*F*(1,11)=5.220, *p*=0.043; **Fig. 4E**). In addition, chemogenetic mPFC disinhibition tended to impair the expression of the old rule prior to rule change, as reflected by a trend toward a reduced percentage of correct responses during the reminder trials (*F*(1,11)=3.783, *p*=0.078) (**Fig. 4F**).

Notably, the serial reversal impairment caused by chemogenetic mPFC disinhibition presented itself without a substantial impact on response latencies or omission rate (**Table S3**). Systemic injection of 4.5 mg/kg of CNO-2HCl only tended to increase latencies following reversal (*F*(1,10)=3.725, *p*=0.082), irrespective of response type (*F*<1), but did not affect response latencies during reminder trials (main effect or interaction involving drug, largest *F*(1,11)=2.292, *p*=0.158). No other effect or interaction was observed with respect to omissions or response latencies (largest *F*(1,11)=2.164, *p*=0.164).

#### Bayesian strategy analysis: chemogenetic mPFC disinhibition induced subtle impairments in task-pertinent strategy implementation around reversal

We also examined the effect of chemogenetic mPFC disinhibition on the probability of task-pertinent strategies around reversal that we had found to be disrupted by mPFC picrotoxin (**Fig. 4G-I**). Chemogenetic mPFC disinhibition numerically reduced adherence to the old rule (go-previous), which was lower following CNO-2HCl (4.5 mg/kg) compared to saline injection across responses around reversal (6 responses before, and 10 after reversal), although this effect was not significant (*F*(1,11)=3.038, *p*=0.109; drug × response interaction, *F*<1) (**Fig. 4G**). Chemogenetic mPFC disinhibition subtly changed lose-shift behaviour: compared to saline, CNO-2HCl reduced lose-shift probability across the 6 responses before reversal but increased it across the 10 responses following reversal; the latter reflected that lose-shift probabilities following reversal declined less markedly in the CNO-2HCl condition compared to saline (**Fig. 4H**). This was supported by a significant drug × response interaction (*F*(15,165)=2.691, *p*=0.001), although simple main effects analysis did not reveal a significant injection effect at any single response (largest *F*(1,11)=1465, *p*=0.254). Finally, chemogenetic mPFC disinhibition tended to reduce win-stay behavior around reversal (**Fig. 4I**). This was supported by a trend towards a main effect of drug (*F*(1,11)=3.537, *p*=0.087), in absence of a drug × response interaction (*F*(15, 165)=1.341, *p*=0.183).

Overall, mPFC disinhibition by chemogenetically inhibiting GABAergic neurons within the mPFC impaired serial reversal learning, replicating our finding following mPFC disinhibition by picrotoxin. Our findings suggest that mPFC disinhibition primarily disrupts serial reversal learning by increasing regressive errors, as this increase was evident both following chemogenetic mPFC disinhibition with 4.5 mg/kg of CNO-2HCl and following mPFC picrotoxin. Moreover, mPFC picrotoxin reduced win-stay behavior and mPFC chemogenetic disinhibition tended to have the same effect, suggesting that reduction in this exploitative behavior is also important for the disruption of reversal learning by mPFC disinhibition. In addition, mPFC picrotoxin also increased perseverative responding, reduced explorative lose-shift behavior and increased response latencies and omissions. These effects do not seem to be required for the serial reversal impairment, because they were not observed following chemogenetic mPFC disinhibition with 4.5 mg/kg CNO-2HCl, although such chemogenetic mPFC disinhibition caused a similarly marked serial-reversal learning impairment as mPFC picrotoxin.

#### CNO-2HCl did not affect serial reversal performance in control rats

Finally, we completed a control experiment to confirm that CNO did not impair serial reversal learning in rats not expressing the DREADD (n=16; **Table 1**, Cohort #5) (data not shown). Comparing serial reversal performance following injection of 4.5 mg/kg CNO-2HCl or of saline, we observed no significant differences in RTC, reminder-trial performance, regressive errors, response latencies, or omissions (largest *F*(1,12)=1.695, p=0.217), and only a trend for CNO-2HCl to increase perseverative errors (*F*(1,15)=3.305, p=0.089). These findings support that CNO-2HCl does not affect serial reversal learning independently of DREADD activation.

## DISCUSSION

During early reversals (R1 to R3), control rats showed a marked ‘reversal cost’ at R1, manifested by increased RTC compared to the SD stage, as well as increased perseverative errors (**Fig. 1C, D**); both RTC and errors numerically declined from R1 to R3, reflecting that rats ‘learned to reverse’. In contrast, during later serial reversals (R5 to R10), after rats had had the opportunity to ‘learn to reverse’, RTC were lower than at the SD stage and there were less errors than during early reversals, including similar numbers of perseverative and regressive errors (**Fig. 2C-E**). mPFC functional inhibition by muscimol impaired early-reversal performance, increasing RTC (**Fig. 1C**) and perseverative errors (**Fig. 1D**) during R2, but did not markedly affect serial-reversal performance (apart from slightly increasing win-stay and sticky behavior during responses leading up to criterion performance). This early-reversal impairment was associated with increased use of the old strategy (go-previous, **Fig. 1G**) and reduced exploration (lose-shift behavior, **Fig. 1H**), but intact exploitation (win-stay, **Fig. 1I**), following reversal. In contrast, mPFC disinhibition by picrotoxin facilitated early reversal performance at R2, likely due to impaired expression of the previous response, and markedly disrupted serial-reversal performance, increasing RTC (**Fig. 2C**) and both perseverative and regressive errors (**Fig. 2D&E**). This serial-reversal impairment by mPFC picrotoxin presented alongside the impaired expression of the old response (reduced percentage of correct reminder-trial responses, **Fig. 2F**). It was associated with impaired win-stay behavior (**Fig. 2I**) and some reduction of lose shift behaviour (**Fig. 2H**), as well as reduced use of the old strategy around reversal (**Fig. 2G**). Next, we used Cre-dependent expression of the inhibitory hM4Di DREADD in the mPFC of Vgat-Cre rats to selectively inhibit mPFC GABAergic neurons. We verified that the DREADD was predominantly expressed in GABAergic neurons and showed that DREADD activation particularly enhanced mPFC burst firing and increased locomotor activity (**Fig. 3**). Importantly, chemogenetic mPFC disinhibition also markedly impaired reversal performance, increasing RTC (**Fig. 4C**) and regressive errors (**Fig. 4E**), and tending to reduce win-stay behavior (**Fig. 4I**), without affecting perseverative errors and lose-shift behaviour after reversal.

### mPFC functional inhibition impaired early, but not serial, reversals

mPFC functional inhibition impaired early reversal learning, specifically at R2, but otherwise left reversal performance intact, confirming and extending previous studies. In similar two-lever reversal procedures in rats, mPFC functional inactivation (by the sodium channel blocker bupivacaine) did not affect performance at R1 (Floresco et al., 2008) and mPFC functional inhibition (by muscimol and baclofen) did not impair later serial-reversal performance (Hervig et al., 2020), but performance at R2 was not examined. Another study, testing the impact of cytotoxic mPFC lesions in rats performing two-odor serial reversals, reported findings strikingly similar to ours, namely a selective impairment at R2, but not other reversal stages (testing was continued up to R8) (Kinoshita et al., 2008). Like mPFC functional inhibition in the present study, mPFC lesions did not impair performance beyond the ’normal’ reversal cost at R1, but impaired rats’ ability to overcome this reversal cost at R2. These findings are consistent with the idea that the mPFC is required to overcome prepotent behavioral responses (Haddon & Killcross, 2007; Marquis et al., 2007; Miller & Cohen, 2001), in this case, pressing the previously correct lever.

In the present study, mPFC functional inhibition impaired reversal at R2 by increasing perseverative errors and decreasing lose-shift behavior. Similarly, reversal impairments by mPFC lesions on a three-lever spatial-discrimination serial-reversal task (Kosaki & Watanabe, 2012) or by mPFC functional inhibition on a plus-maze serial-spatial reversal task (Avigan et al., 2020) mainly reflected increased perseverative errors. Moreover, similar to the present study, reduced lose-shift behavior was reported following mPFC functional inactivation on a probabilistic two-lever spatial-reversal task (Dalton et al., 2016), and following mPFC lesions on bowl-digging set-shift or reversal tasks requiring olfactory and tactile discriminations (Wang et al., 2019). Interestingly, on the probabilistic two-lever spatial-reversal task, where only 80% of correct choices were rewarded, the reduced lose-shift behavior was associated with better reversal performance (Dalton et al., 2016), possibly reflecting that it was advantageous for rats to shift less readily following a non-rewarded response. In contrast, in the present study and on the bowl-digging set-shift and reversal tasks (Wang et al., 2019), where responses were deterministic reduced lose-shift behavior was associated with impaired reversal or set-shift performance. These findings support that the mPFC can contribute to reversal performance by suppressing perseverative errors and sustaining lose-shift behavior. On tasks with deterministic reward contingencies, mPFC-dependent lose-shift behavior facilitates reversal performance (present study; Wang et al., 2019), whereas, on tasks with probabilistic reward contingencies, such lose-shift behavior may hinder reversal performance (Dalton et al., 2016).

The mPFC may contribute to reversal learning by other cognitive mechanisms. First, mPFC lesions impaired a touch-screen visual-pattern reversal task mainly by increasing regressive errors and only when difficult to discriminate visual stimuli were used; this was suggested to reflect that the mPFC is important for visual attention (Brigman & Rothblat, 2008; Bussey et al., 1997; Chudasama & Robbins, 2003), and may also reflect the importance of the mPFC for attentional sets (Birrell & Brown, 2000; Brown & Bowman, 2002). However, attentional impairment is unlikely to explain the reversal-learning impairments caused by mPFC functional inhibition in the present study. This is because mPFC functional inhibition selectively increased perseverative errors, whereas an increase in regressive errors would be expected on the basis of an attentional impairment (Brigman & Rothblat, 2008; Bussey et al., 1997); moreover, although both mPFC functional inhibition and disinhibition impaired attention (Auger et al., 2017; Pezze et al., 2014), only functional inhibition impaired early reversal learning. Second, some studies reported that the mPFC was required for serial spatial-reversal learning on plus-maze tasks (but see: Rich & Shapiro, 2007; Young & Shapiro, 2009), and this was suggested to reflect that the mPFC is required for working memory (Kidder et al., 2024) and for reducing proactive interference based on hippocampus-dependent spatial memory (Avigan et al., 2020; Guise & Shapiro, 2017). These cognitive mechanisms may also contribute to performance on our two-lever reversal task. However, disruption of these processes cannot easily account for the specific impairments at R2 following mPFC functional inhibition.

### mPFC disinhibition impaired serial, but not early, reversals

mPFC disinhibition, by the GABA-A receptor antagonist picrotoxin or chemogenetic inhibition of GABAergic neurons, markedly impaired serial reversal learning. Both types of mPFC disinhibition increased RTC and regressive errors and tended to decrease win-stay behavior. This suggests that a key effect of mPFC disinhibition is impaired exploitation, i.e. selection of rewarded behaviors. In addition, mPFC picrotoxin impaired exploration, reflected by reduced lose-shift behaviour and increased perseverative errors, and increased response latencies and omissions. These additional effects of mPFC picrotoxin were not critical for the serial-reversal learning impairment, which was similarly pronounced following mPFC picrotoxin or chemogenetic disinhibition.

Contrasting with the marked serial reversal impairments, mPFC disinhibition by picrotoxin did not disrupt early reversals, consistent with previous work (Enomoto et al., 2011), and even facilitated reversals at R2. The latter was at least partly due to a high number of omissions, which limited reminder-trial responses and reduced reinforcement of the old rule, rather than improved behavioral flexibility.

Electrophysiological recordings showed that mPFC disinhibition intensifies burst firing, whereas functional inhibition reduced firing, including burst firing, of mPFC neurons (Pezze et al., 2014; present study). Only ‘too little’ mPFC activity caused by functional inhibition, but not ‘too much’ mPFC activity caused by disinhibition disrupted early reversals, specifically at R2, whereas the opposite was true for serial reversals. In contrast, both mPFC functional inhibition and disinhibition impaired attention (Auger et al., 2017; Pezze et al., 2014). This highlights that distinct cognitive functions can show distinct relationships to mPFC activity, and that mPFC disinhibition can disrupt functions that do not require the disinhibited region (i.e., are not disrupted by regional functional inhibition or lesion), presumably because disinhibition disrupts processing in mPFC projection sites (Bast et al., 2017).

Although the downstream effects of mPFC disinhibition remain to be investigated, several mPFC projection sites may mediate the serial-reversal impairments by mPFC disinhibition, including the OFC (Sesack et al., 1989), dorsal and ventral striatum (Mailly et al., 2013) and lateral habenula (Baker et al., 2015). The OFC, especially lateral OFC, is required for serial-reversal learning, to maintain appropriate exploration and exploitation (Hervig et al., 2020; Murray et al., 2007). The striatum is also required for reversal learning, with dorsomedial striatal lesions impairing serial reversals by increasing perseveration (Castañé et al., 2010; Clarke et al., 2008; Young et al., 2023), whereas ventral striatal (nucleus accumbens shell) inactivation impairing serial reversals by reducing win-stay behavior (Dalton et al., 2014). Finally, the habenula has been implicated in behavioral flexibility; lateral habenula functional inhibition impaired serial T-maze reversals, increasing regressive errors, reducing win-stay behavior and increasing lose-shift behavior (Baker et al., 2015).

### Conclusion and clinical implications

Our findings suggest that reduced mPFC activation can impair early reversal learning by disrupting exploration necessary to overcome the early-reversal cost, whereas mPFC disinhibition can disrupt serial reversals, especially by disrupting exploitation, possibly due to aberrant drive of mPFC projection sites. Considering the functional-anatomical similarities between rodent mPFC and primate dlPFC (see *Introduction*), our findings suggest that dlPFC disinhibition and hypofrontality could both contribute to reversal-learning deficits, which characterize many neuropsychiatric disorders (Izquierdo et al., 2017). This is particularly relevant to schizophrenia, which has been associated with both dlPFC disinhibition and hypofrontality (see *Introduction*).

## Author contributions

JR, MvH, SD, JDH, PMM, CWS, SM, and TB designed research.

JR, CT, LOH, RGA, JL, JJ performed research.

JR and SM analyzed data.

JR and TB wrote the original draft of the paper, with all other authors subsequently contributing to revising the draft; all authors have approved the final content.

## Acknowledgements

We thank the staff of the Biological Support Unit (BSU) for their contributions to the husbandry and welfare of the rats used in the present study. In addition, we thank Matthias Heil (B.I.) and the School of Life Science Imaging facilities (SLIM; University of Nottingham) for their guidance and expertise on histological and tissue imaging procedures included in this report.

