## Supplemental Figures for "Too little and too much: medial prefrontal functional inhibition impairs early, whereas neural disinhibition impairs serial reversal performance in rats"

### Supplementary tables and figures

**mPFC saline/picrotoxin/muscimol, early reversal learning**

| **Reversal stage** | **Saline (0.9%)** | | | **Picrotoxin (300ng)** | | | **Muscimol (62.5ng)** | | |
|---|---|---|---|---|---|---|---|---|---|
| | Omissions | $Lat_{Cor}$ (s) | $Lat_{Inc}$ (s) | Omissions | $Lat_{Cor}$ (s) | $Lat_{Inc}$ (s) | Omissions | $Lat_{Cor}$ (s) | $Lat_{Inc}$ (s) |
| | *n=15* | *n=15* | *n=15* | *n=14* | *n=12* | *n=12* | *n=15* | *n=15* | *n=15* |
| SD | 9.00±5.66 | 0.99±0.07 | 1.31±0.09 | 108.92±17.91 | 2.64±0.18 | 2.94±0.19 | 2.40±0.84 | 1.17±0.09 | 1.32±0.10 |
| R1 | (0.33±0.16)<br>1.13±0.36 | (1.08±0.14)<br>0.94±0.10 | (1.15±0.23)<br>1.10±0.11 | (14.5±1.41)<br>65.14±13.15 | 1.98±0.25 | 2.75±0.21 | (0.53±0.19)<br>3.20±0.86 | (1.18±0.12)<br>1.15±0.09 | (1.75±0.23)<br>1.24±0.12 |
| R2 | (0.07±0.07)<br>0.53±0.17 | (0.99±0.15)<br>1.07±0.11 | (0.96±0.09)<br>1.06±0.11 | (17.7±1.22)<br>58.36±7.65 | 2.24±0.27 | 1.92±0.12 | (0.13±0.09)<br>2.80±0.67 | (1.06±0.17)<br>1.20±0.11 | (1.08±0.10)<br>1.26±0.08 |
| R3 | (0.27±0.21)<br>1.07±0.55 | (0.94±0.10)<br>0.96±0.10 | (1.03±0.19)<br>0.99±0.10 | 15.6±1.94)<br>49.50±11.24 | 1.65±0.19 | 1.81±0.23 | (0.33±0.27)<br>1.87±0.92 | (1.20±0.16)<br>1.20±0.13 | (1.53±0.32)<br>1.57±0.22 |

**Table S1.** *Omissions and response latencies for early reversal learning experiment comparing impact of mPFC saline, picrotoxin and muscimol.* Values (mean±SEM) are shown for the different task stages (spatial discrimination, SD; reversals 1 to 3, R1 to 3). Values during the 20 reminder trials at the beginning of each reversal stage are shown in brackets. Response latencies are indicated separately for correct responses ($Lat_{Cor}$) and incorrect responses ($Lat_{Inc}$). Sample sizes contributing to the mean values are indicated at the top of each column. Note, only one rat in the picrotoxin group completed at least one correct and one incorrect response and, therefore, this group was excluded from the analysis of reminder-trial response latencies and no reminder trial response latencies are shown for this group. Moreover, two rats in the picrotoxin group did not make at least one incorrect response across all task stages and had to be excluded completely from the latency analysis, resulting in a reduced sample size of n=12 (instead of 14) for the latency means.

**mPFC saline/picrotoxin/muscimol, serial reversal learning**

| Infusion series | Saline (0.9%) | | | Picrotoxin (300ng) | | | Muscimol (62.5ng) | | |
|---|---|---|---|---|---|---|---|---|---|
| | Omissions | $Lat_{Cor}$ (s) | $Lat_{Inc}$ (s) | Omissions | $Lat_{Cor}$ (s) | $Lat_{Inc}$ (s) | Omissions | $Lat_{Cor}$ (s) | $Lat_{Inc}$ (s) |
| | *n=13* | *(n=12)*<br>*n=13* | *(n=12)*<br>*n=13* | *n=13* | *(n=12)*<br>*n=13* | *(n=12)*<br>*n=13* | *n=13* | *(n=12)*<br>*n=13* | *(n=12)*<br>*n=13* |
| Series 1 | (0.15±0.10)<br>0.46±0.18 | (0.66±0.08)<br>0.59±0.05 | (0.87±0.17)<br>0.59±0.05 | (13.62±6.77)<br>34.77±8.29 | (2.65±0.27)<br>2.37±0.35 | (2.42±0.15)<br>2.99±0.39 | (0.23±0.17)<br>1.00±0.36 | (0.67±0.10)<br>0.66±0.10 | (0.76±0.14)<br>0.86±0.14 |
| Series 2 | (0.31±0.21)<br>0.38±0.21 | (0.65±0.09)<br>0.50±0.04 | (0.70±0.19)<br>0.66±0.10 | (2.77±0.79)<br>23.54±7.64 | (2.77±0.36)<br>1.44±0.18 | (2.39±0.44)<br>1.95±0.21 | (0.15±0.10)<br>0.31±0.13 | (0.59±0.08)<br>0.69±0.06 | (1.10±0.27)<br>0.63±0.08 |

**Table S2.** *Omissions and response latencies for serial reversal learning experiment comparing impact of mPFC, saline, picrotoxin and muscimol.* Values (mean±SEM) are shown for the two-infusion series (series 1, reversals 5 to 7, R5 to 7; series 2, reversals 8-10, R8 to 10). Values during the 20 reminder trials at the beginning of each reversal stage are shown in brackets. Response latencies are indicated separately for correct responses ($Lat_{Cor}$) and incorrect responses ($Lat_{Inc}$). Sample sizes contributing to the mean values are indicated at the top of each column. Note: one rat did not make at least one incorrect response during reminder trials in all three drug conditions in both series and, therefore, was excluded from the analysis of reminder trial response latencies; the resulting sample size for $Lat_{Inc}$ during reminder trials is indicated in brackets.

**Chemogenetic disinhibition, serial reversal learning**

| Vehicle (saline, 0.9%) | | | CNO-2HCl (4.5mg/kg) | | |
|---|---|---|---|---|---|
| Omissions | $Lat_{Cor}$ (s) | $Lat_{Inc}$ (s) | Omissions | $Lat_{Cor}$ (s) | $Lat_{Inc}$ (s) |
| *n=12* | *n=12* | *n=12* | *n=12* | *n=12* | *n=12* |
| (0±0) | (0.70±0.13) | (0.53±0.07) | (1.58±1.08) | (0.90±0.24) | (0.86±0.20) |
| 0.16±0.11 | 0.63±0.10 | 0.70±0.11 | 4.83±3.85 | 1.00±0.18 | 1.22±0.26 |

**Table S3.** *Omissions and response latencies for serial reversal learning experiment comparing the impact of chemogenetic mPFC disinhibition by 4.5 mg/kg CNO-2HCl to saline injection.* Values are shown as mean±SEM. Values during the 20 reminder trials at the beginning of each reversal stage are shown in brackets. Response latencies are indicated separately for correct responses ($Lat_{Cor}$) and incorrect responses ($Lat_{Inc}$). Sample sizes contributing to the mean values are indicated at the top of each column.

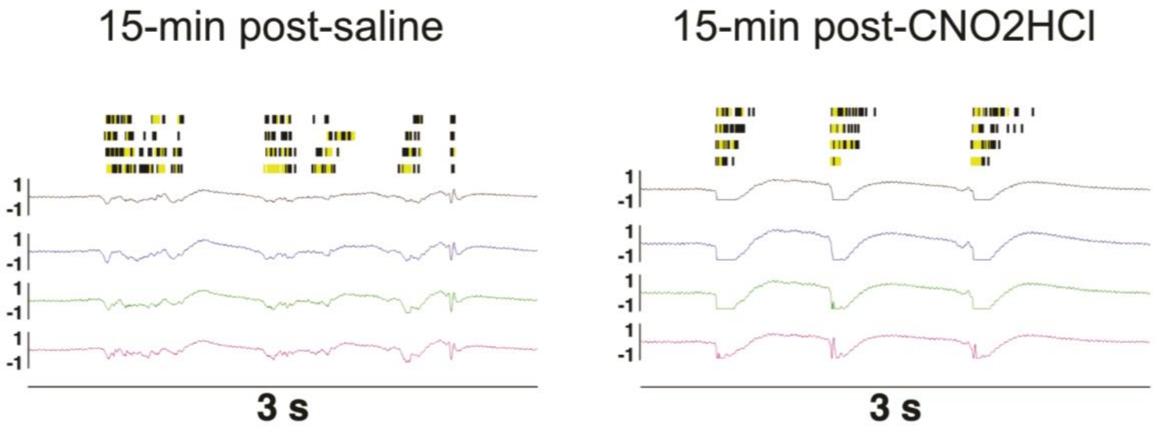


**Figure S1.** *Chemogenetic disinhibition following 6.0 mg/kg CNO2HC enhanced LFP spike-wave discharges and intensified burst firing within the mPFC.* Raster plot (top, black vertical lines; yellow shaded areas indicate automatically classified 'bursts') and LFP traces (bottom, colored lines) for one electrode array (electrodes 1-4) during a 3 second period 15-min post-saline (left) or 6.0 mg/kg CNO-2HCl (right). Following mPFC CNO-2HCl, there was a notable increase in potent LFP spike-wave discharges (negative deflection followed by a positive wave), alongside intensification of multiunit burst firing. These changes in mPFC neural activity were similar to those caused mPFC disinhibition by picrotoxin microinfusion in our previous study, including spike-wave LFP discharges and enhanced LFP power and enhanced neural burst firing; Pezze et al., 2014). Note, negative LFP spikes following CNO-2HCl were clipped, because they exceeded our LFP amplitude recording range.

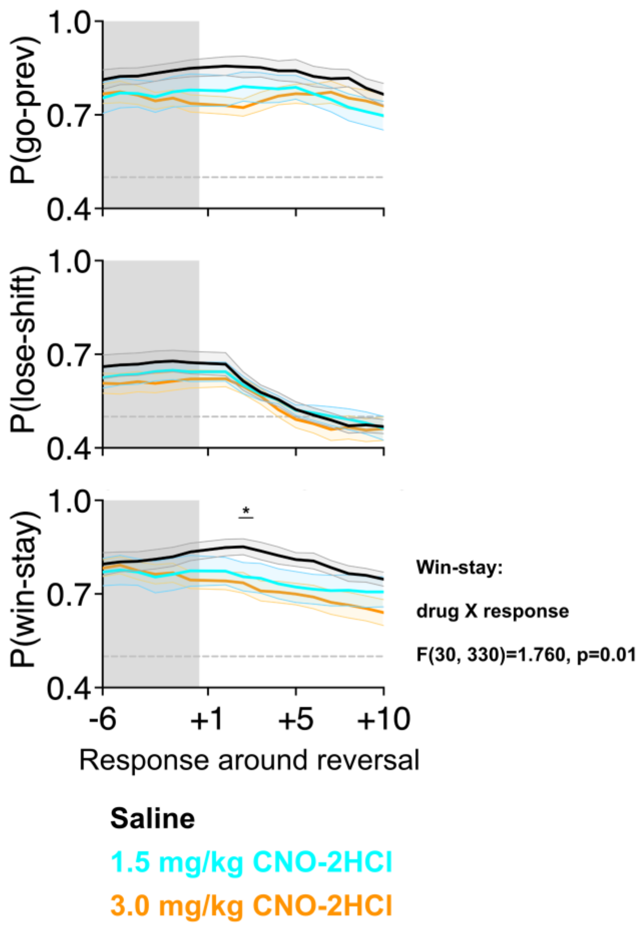


**Figure S2.** *Response strategies around reversal following chemogenetic mPFC disinhibition by 1.5 (light blue) and 3.0 mg/kg (gold) CNO-2HCl, compared tosaline (black) injection*. (Go-previous [top], lose-shift [middle], and win-stay [bottom]). Coloured lines and shaded regions indicate mean±SEM. Grey-shaded region indicates reminder trials. Asterisks above datapoints indicate responses where groups significantly differed ($p<0.05$) following observation of a drug × response interaction. Details on significant interaction included in the figure.

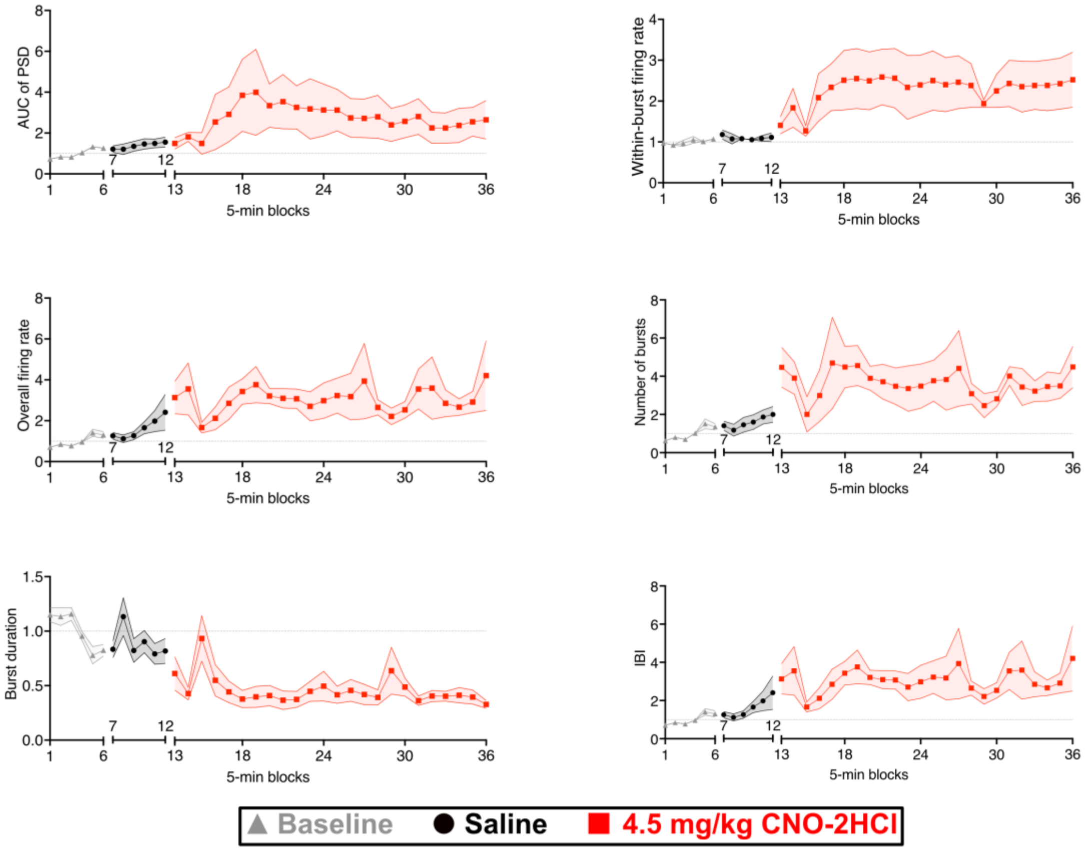


**Figure S3.** *Enhancement of LFP power and multi-unit burst firing by chemogenetic mPFC disinhibition: time course of LFP and multi-unit parameter across baseline (grey triangles) and following systemic saline (black circles) and 4.5 mg/kg CNO-2HCl (red squares) injection.* A) AUC of PSD, B) within-burst firing rate, C) overall firing rate D) number of bursts per block, D) burst duration F) inter-burst interval. All testing was run within-subjects with saline injection always preceding CNO-2HCl. Shaded regions x̄±SEM, and all values were normalized to baseline prior to averaging across rats.
